# A 3D gene expression atlas of the floral meristem based on spatial reconstruction of single nucleus RNA sequencing data

**DOI:** 10.1101/2021.06.30.450319

**Authors:** Manuel Neumann, Xiaocai Xu, Cezary Smaczniak, Julia Schumacher, Wenhao Yan, Nils Blüthgen, Thomas Greb, Henrik Jönsson, Jan Traas, Kerstin Kaufmann, Jose M Muino

**Author notes:** Joint Authors (Manuel Neumann, Xiaocai Xu).

## Abstract

Identity and functions of plant cells are influenced by their precise cellular location within the plant body. Cellular heterogeneity in growth and differentiation trajectories results in organ patterning. Therefore, assessing this heterogeneity at molecular scale is a major question in developmental biology. Single-cell transcriptomics (scRNA-seq) allows to characterize and quantify gene expression heterogeneity in developing organs at unprecedented resolution. However, the original physical location of the cell is lost during the scRNA-seq procedure. To recover the original location of cells is essential to link gene activity with cellular function and morphology. Here, we reconstruct genome-wide gene expression patterns of individual cells in a floral meristem by combining single-nuclei RNA-seq with 3D spatial reconstruction. By this, gene expression differences among meristematic domains giving rise to different tissue and organ types can be determined. As a proof of principle, the data are used to trace the initiation of vascular identity within the floral meristem. Our work demonstrates the power of spatially reconstructed single cell transcriptome atlases to understand plant morphogenesis. The floral meristem 3D gene expression atlas can be accessed at http://threed-flower-meristem.herokuapp.com

## INTRODUCTION

Characterizing gene expression dynamics and heterogeneity at single cell resolution is essential to understand the molecular mechanisms underlying cellular differentiation in multicellular organisms. Technologies based on cell dissociation (e.g. Denyer et al. 2019) or nuclei isolation (e.g. Sunaga-Franze et al. 2021) combined with high-throughput transcriptome sequencing (scRNA-seq/snRNA-seq) allow to characterize the transcriptomes of hundreds of thousands cells at single-cell resolution. However, the physical location of these cells is lost during the experimental process. In plants and other multicellular organisms, cell fate strongly depends on its precise position within the developing organism (Xu et al. 2021). Therefore, it is essential to characterize gene expression patterns of each cell in their native physical context to fully understand the link between gene activity and organ development.

In recent years, there has been a strong development in the field of spatial transcriptomics (Marx 2021; Waylen et al. 2020). However, to date, only one study in plants has been published using an early version of the 10x Visium technology with limited cellular resolution (Giacomello et al. 2017). This lack of technological adaptation of spatial transcriptomics to plants may be because of the difficulties with the enzymatic permeabilization of the cell wall. Single molecule FISH (smFISH) and other high-resolution FISH experiments are also rarely used in plant studies (e.g. Duncan et al. 2016; Solanki et al. 2020) due to the endogenous autofluorescence of many plant cells (Duncan et al. 2016).

Computational inference of spatial locations of cells by mapping scRNA-seq transcriptomes into a computationally binned representation of the studied structure provides an alternative for spatial reconstruction of omics data. In an early study, spatial reconstruction was performed by integrating the expression of nearly 100 reference genes using a mixture model (Stuart et al. 2019). Allowing better resolution, *DistMap* (Karaiskos et al. 2017) assigns scRNA-seq cells to individual cells of a computationally generated spatial map containing the expression of ∼80 reference genes using Matthews correlation coefficient. Other methods aim to combine scRNA-seq with high-throughput spatial maps (e.g. MERFISH, Slide-seq) that collect the expression of thousands of reference genes. They are based on the projection of the scRNA-seq and the spatial map transcriptomes into a common latent space (SEURAT (Satija et al. 2015), Liger (Welch et al. 2019), Harmony (Korsunsky et al. 2019), gimVI (Lopez et al. 2018), SpaGe (Abdelaal et al. 2020)). In general, there is a tendency to develop computational methods that require a large number of reference genes, which limits these tools to organisms with extensive spatial transcriptomics resources.

In plants, spatio-temporal gene expression patterns are usually established using traditional *in situ hybridization* or by confocal microscopy of promoter fusions to fluorescent reporters. Confocal microscopy has the advantage that it can be used to reconstruct 3D structures by combining several z-stack images (Vijayan et al. 2021; Hernandez-Lagana et al. 2021; Wolny et al. 2020; Bravo González-Blas et al. 2020; Refahi et al. 2021). In addition, combined with live image microscopy the temporal dynamics of gene expression and morphology development can be reconstructed (Refahi et al. 2021; Valuchova et al. 2020). In this way, Refahi et al. (2021) combined the information on spatio-temporal expression patterns of 28 regulatory genes into 3D reconstructed *Arabidopsis* flower meristems, ranging from initiation to stage 4 of flower development. These methods are limited by the low number of genes profiled per experiment, Therefore, tools to integrate scRNA-seq with expression data of defined, limited sets of 3D reference gene expression patterns need to be developed for spatial reconstruction single cell transcriptomes in plants.

Here, we adapted novoSpaRc (Nitzan et al. 2019), a methodology for spatial reconstruction of single cell RNA-seq data, to generate a 3D single-cell transcriptome atlas of a floral meristem by integrating single nuclei RNA-seq and a 3D reconstructed flower meristem (Refahi et al. 2021). NovoSpaRc reconstruction aims to explicitly preserve the transcriptome similarity among closely located scRNA-seq cells in the spatial map, while maximizing the transcriptome similarity between the scRNA-seq cells and the cells of the spatial map to which they are assigned. In such a way, novoSpaRc performance is less affected by the number of reference genes than other methods, and, in theory, it can also be used without any reference gene (Nitzan et al. 2019). However, novoSpaRc was developed to make use of spatial 2D continuous reference gene expression maps, while the 3D expression spatial map of floral meristem generated by Refahi et al. 2021 is binary. We adapted the methodology for reconstructing single-cell transcriptomes in 3D making use of binary reference gene expression data. By this, we were able to generate an atlas of gene expression in different meristematic domains and spatially trace the earliest stages of tissue differentiation within the *Arabidopsis* flower. In summary, these results provide a primer for future initiatives to generate plant organ 3D atlases of gene expression.

## RESULTS

### snRNA-seq of *Arabidopsis* floral meristems

In order to obtain genome-wide gene expression profiles in the floral meristem at single cell level, we use a system for synchronized floral induction (*pAP1*:AP1-GR *ap1-1 cal-1*; (Kaufmann et al. 2010) to maximize the collection of plant material from the desired developmental stage. We chose to study stage 5 of flower development because of the availability of several –omics datasets from this stage (e.g. Pajoro et al. 2014; Wuest et al. 2012; Ó’Maoiléidigh et al. 2013), which are needed to validate the performance of the method. At stage 4-5 (Smyth, Bowman, and Meyerowitz 1990), the whorled organization of the flower gets established, and homeotic gene activity defines domains within the meristem that will give rise to different floral organ types, therefore being an excellent stage to study the initial steps of floral organ specification.

We performed single nuclei RNA-seq (snRNA-seq), where nuclei were collected by fluorescence-activated DAPI-stained nuclei sorting (FANS), and snRNA-seq datasets were created using the 10x Chromium system. In this way, Cell Ranger v3.1.0 identified 7,716 single nuclei transcriptomes with a median of 1,110 genes expressed per nucleus. The low number of reads mapping to mitochondria genes (<5%) indicates a low organelle contamination (Sup Fig 1). Fig 1A shows that snRNA-seq is able to recapitulate (R=0.88) the expression profile of bulk RNA-seq data obtained from the same stage and tissue type. Analysis of the data using Seurat v3.2.3 identified 12 main clusters and the marker genes defining these clusters (Sup Table 1). To annotate the clusters, we identified the top 20 marker genes specific for each cluster and we plotted the expression of these marker genes in publically available bulk RNA-seq datasets of different tissues and floral stages (Fig 1D, and Sup Fig 2). In addition, we calculated the average expression of known floral meristem marker genes in the 12 snRNA-seq clusters (Fig 1C).

**Figure 1.**
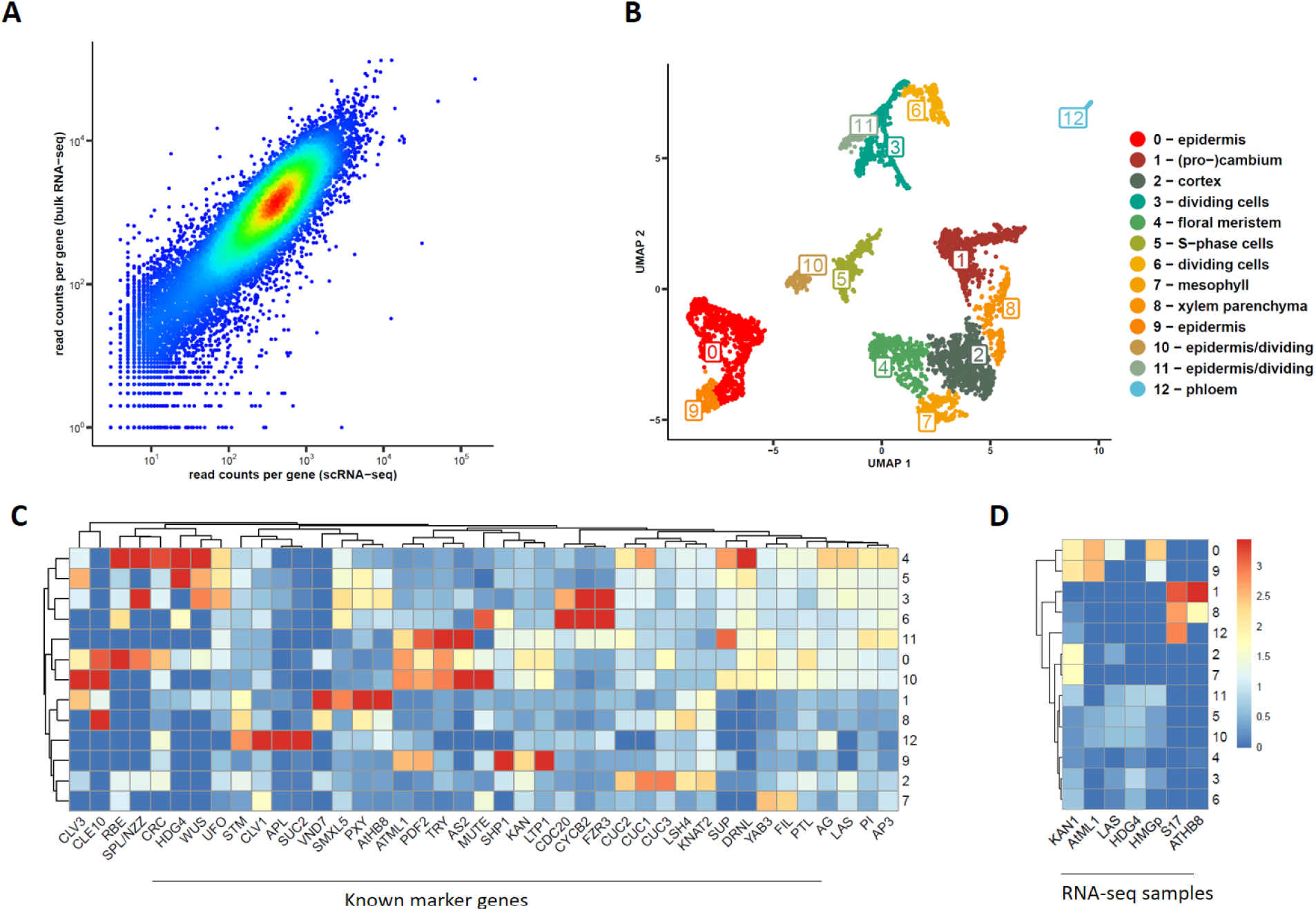
Single-nucleus RNA-sequencing of Arabidopsis floral meristems. A) Reproducibility (R= 0.88) of the gene expression estimated from computationally pooling all nuclei from our snRNA-seq compared to bulk-RNA-seq of stage 5 flower meristem (average of 3 biological replicates). B) UMAP plot and clustering snRNA-seq analysis of Arabidopsis floral meristems obtained by Seurat analysis. C) Average expression of known floral markers on the identified snRNA-seq clusters. D) Average expression of the top 20 marker genes for each snRNA-seq cluster (y-axis) on domain-specific shoot apical meristem bulk RNA-seq datasets profiled by (Tian et al. 2019) (x-axis). See Sup. Figure 2 for expression profiles in other plant domains/stages.

We were able to recover the main tissue types present in the meristem, including different epidermal as well as vascular tissue types. The four epidermis clusters (0, 9, 10 and 11) show specific expression of *MERISTEM LAYER 1 (ATML1)* (Sessions, Weigel, and Yanofsky 1999) and *PROTODERMAL FACTOR 1/2 (PDF1/2)* (Abe et al. 2003) (Sup Table 1, Sup Fig 3). Clusters 0 and 9 are distinguished by the expression of individual marker genes such as *TRIPTYCHON (TRY)* (Pesch and Hülskamp 2011)*, TRICHOMELESS1 (TCL1)* (Wang et al. 2007), and genes involved in wax composition which indicates epidermal cells that will develop trichomes (cluster 0) or not (cluster 9). Cluster 10 and 11 represent dividing epidermal cells, marked by the expression of genes coding for histones which is characteristic of the S-phase and genes involved in cell division (Sup Table 1, Sup Fig 3).

Clusters 1, 8 and 12 can be classified as vasculature (Fig. 1D). More specifically, cluster 1 corresponds to vascular stem cells, as marked by cambium (Sup Fig 2C) expressing markers genes such as *PHLOEM INTERCALATED WITH XYLEM (PXY)* and *SMAX1-LIKE 5 (SMXL5)* (Shi et al. 2021) (Sup Fig 3). Cluster 12 contains cells that are associated with phloem, containing the marker genes *ALTERED PHLOEM DEVELOPMENT* (*APL*) (Shi et al. 2021; Bonke et al. 2003) (Sup Fig 3). Cluster 8 is enriched for vascular xylem parenchyma genes, for example *CYTOCHROME P450, FAMILY 708 (CYP708A3)* (Shi et al. 2021) (Sup Fig 3), and shows signatures of cell expansion and cellulose biosynthesis. It should be noted that in this dataset, no mature xylem vessels or phloem sieve elements can be expected, because these structures lack a nucleus.

The analysis of marker genes of cluster 2 shows an enrichment on genes involved in starch catabolic process as well as genes expressed in the cortex such as *CHALLAH (CHAL)* (Uchida et al. 2012; Sup Fig 3) and *JACKDAW* (*JKD*) (Hassan, Scheres, and Blilou 2010), which indicates that cluster 2 represents cortex . Cluster 4 represents the floral meristem, containing specific markers such as *APETALA3 (AP3)* (Jack, Brockman, and Meyerowitz 1992), *REPRODUCTIVE MERISTEM 34 (REM34)* (Mantegazza et al. 2014) (Sup Fig. 3). Cluster 7 corresponds to cells that differentiate into mesophyll, e.g. in sepals or pedicel, and it shows specific expression of marker genes such as *LIPOXYGENASE 2 (LOX2)* (Jensen, Raventos, and Mundy 2002) (Sup Fig 3) and *REDUCED CHLOROPLAST COVERAGE (REC1)* (Larkin et al. 2016).

Clusters 3, 5, 6, 10, and 11 denote dividing cells (Sup Fig 3). Cluster 3 is a cluster showing enriched expression of several cell-cycle associated genes. Cluster 5 shows specific activation of many histone genes whose activity is associated with the S-phase of the cell cycle, as well as some genes involved in cell proliferation and cell growth (e.g. *AINTEGUMENTA* (Mizukami and Fischer 2000)). Cluster 6 is enriched in cell cycle markers, in particular *CELL DIVISION CYCLE 20.2 (CDC20.2)* which accumulates in the nucleus from prophase until cytokinesis (Yang, Wightman, and Meyerowitz 2017). Cluster 10 and 11 are epidermal cells in different cell cycle phases as described before.

One of the major drawbacks of this unsupervised clustering approach is that it identifies groups of cells depending on their transcriptome variance, and therefore it may miss cell types of biological interest without sufficient biological variance in the system. For example, we were not able to distinguish clusters representing individual floral whorls, likely because the transcriptome variance between tissue types such as epidermis and vasculature is much greater than between different whorls, at least at this stage of development. In addition, the correspondence of each cell cluster to a particular homogeneous physiological cell type is not guaranteed. For example, cluster 1 represents vascular (pro)cambium, but close inspection of this cluster (Sup Fig 3) reveals specific expression of *PXY* (marker of proximal cambium) and *SMXL5* (marker of distal cambium) in separate regions of the cluster. This provides additional justification for the development of a method to map the snRNA-seq transcriptomes to a physical representation of the plant tissue/organ at study. In the next sections, we describe how we map snRNA-seq data to a spatial expression map of the floral meristem that will enable the selection of the group of cells-of-interest (e.g. floral whorls).

### Reconstructing gene expression by snRNA-seq and microscopy image integration

We used novoSpaRc (Nitzan et al. 2019) to integrate snRNA-seq data and a published 3D reconstructed *Arabidopsis* stage 4 floral meristem (“spatial map”) that has information on the expression pattern of 28 genes (“reference genes”) (Refahi et al. 2021). To adapt novoSpaRc to map single nuclei transcriptomes to the 3D floral meristem map with binary expression of the reference genes, we implemented three main modifications:

1. **Filtering:** snRNA-seq was performed on the with ‘cauliflower-like’ meristem plant material, which may contain cells from regions (e.g. short pedicels and stems) that were not present in our spatial map. Therefore, we set up a filtering procedure to eliminate snRNA-seq transcriptomes that were too dis-similar to the transcriptomes of the spatial map (see Material and Methods for details).
2. **Genes used for calculating snRNA-seq transcriptome distances:** The original novoSpaRc pipeline calculates the distance between snRNA-seq transcriptomes using a set of genes selected depending on their variability across the snRNA-seq transcriptome (highly variable genes). Because in our dataset these highly variable genes were not enriched among the known flower marker genes, we also used the top 100 genes with the highest expression correlation with the reference genes, which included very well-known floral regulator genes, to calculate this distance.
3. **Distance used to calculate dissimilarity between spatial map and snRNA-seq transcriptomes:** The original novoSpaRc pipeline calculates distances between transcriptomes from the spatial map and snRNA-seq data using the Euclidean distance. Because our spatial map data is binary, we also employed two other distances commonly used for binary data: Hamming and Jaccard distances.

Subsequently, we studied the performance of these modifications by calculating the area under the receiver operating characteristic (AUROC) for predicting the expression of each reference gene when this gene was removed from the spatial map during the data integration step. Sup. Figure 4 shows the general good performance (AUROC) of our method for each gene and parameter combination tested. Three genes, *HISTIDINE PHOSPHOTRANSFER PROTEIN 6* (*AHP6*), *AUXIN RESPONSE TRANSCRIPTION FACTOR 3* (*ARF3*, *ETTIN*) and *CLAVATA3* (*CLV3*), had very low performance independently of the parameters used (see next paragraph for an explanation). Therefore, we calculated the overall performance of the method as the average AUROC of all genes except *AHP6*, *ETTIN*, *CLV3* and *WUSCHEL (WUS)*. *WUS* was excluded due to the low number of cells (n=8) where it was expressed in the spatial map. In general, modifications improved the performance of the original novoSpaRc pipeline (Sup Fig 5). In particular, using the Jaccard distance had a positive impact on the performance of the method in this particular dataset (Sup Fig. 5). In our hands, other datasets show different optimal parameter settings, but filtering always improves the performance. For visual comparison, Figure 2 shows the reconstructed expression of representative genes when our modifications are applied or not. In particular, *APETALA3* (*AP3*) and *SEPALATA3* (*SEP3*) are the genes showing the biggest differences (see also Sup Fig 4). For the final prediction, modifications and the parameter values which maximized the average AUROC were used to reconstruct gene expression using the whole spatial map dataset (see Material and Methods).

**Figure 2.**
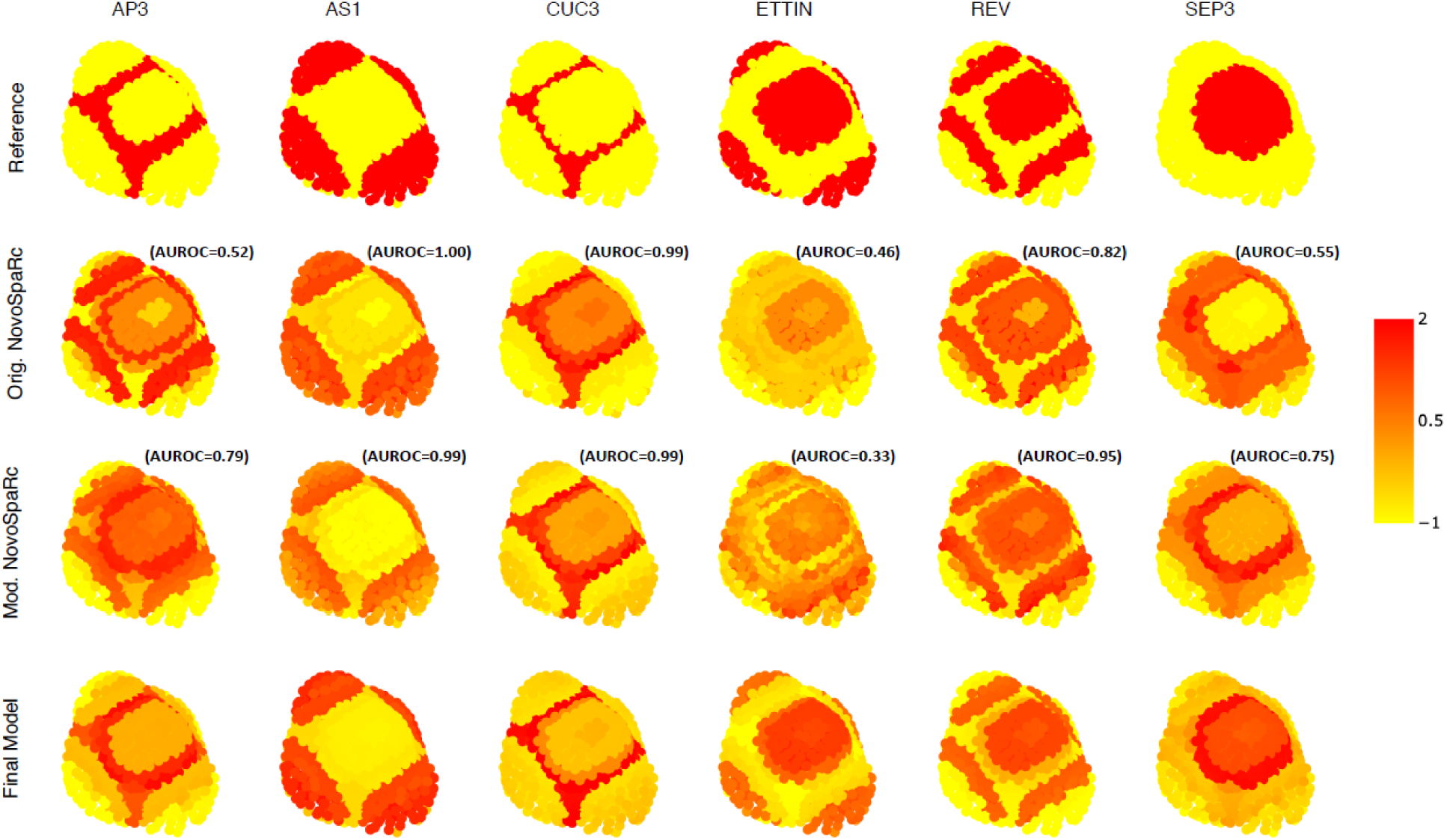
Examples of reconstructed expression patterns for representative genes in Arabidopsis floral meristem. Top row shows the reference expression obtained from the reference spatial map. Second and third row is the reconstructed expression using the parameters that maximize the average AUROC when the gene to be predicted is removed from the data integration step and the original novoSpaRc pipeline (second row) or our modified pipeline (third row) is used. Bottom row is the final reconstructed expression using all the spatial map data. To facilitate visual comparison, we standardized the expression of each gene to have mean 0 and variance 1. The expression of other genes can be visualized at http://threed-flower-meristem.herokuapp.com

As mentioned before, three genes (*ETTIN*, *AHP6* and *CLV3*) had low performance (AUROC close to 0.5) for any set of parameter values used when these genes were removed from the spatial map during the data integration step. We hypothesized that this is because cells expressing these genes are not expressing any of the other reference genes used, and therefore, these cells cannot be correctly mapped. We measured this expression-dependency as the maximum Spearman correlation value of a particular gene against any other gene from the reference list in the snRNA-seq data. We call this value the Predicted Estimation Performance (PEP) for a particular gene. Indeed, there is a strong correlation between the performance of the method (AUROC) and PEP for each gene (Sup Fig 6A), which indicates that we can use it as a predictor of the performance of the method for each particular gene. As we sequentially eliminate genes from the spatial map prior to gene expression reconstruction, starting with the highest correlated reference gene, and therefore decreasing the PEP value of that reconstructed gene, we see a drop in the performance (AUROC) (Sup Fig 6B). However, when we sequentially eliminate reference genes starting with the lowest correlated reference gene, there is no evident decrease of performance (Sup Fig 6C).

Based on Sup Fig 6A, we chose a PEP threshold of 0.13 to decide which genes (n=1,306) we consider to have a reliable expression prediction. We obtained this threshold as the point in Sup Fig 6A where the AUCROC starts to be bigger than 0.5. As the PEP value is estimated without using the spatial map, it can be used to select a set of reference genes for future experiments in order to maximize the number of correctly predicted genes. The number of genes with high PEP values (n=1,306 for PEP>0.13) is mainly influenced by the number of reference genes in the spatial map. Therefore, when using a higher number of reference genes higher PEP values are obtained per gene (Sup Fig 7).

To validate the predictions of spatial gene activity in the floral meristem, we analyzed expression patterns of a set of genes by reporter gene analysis *in planta* (Fig 3). In brief, promoter-GFP fusions were stably expressed in *A. thaliana* and stage 4-5 floral meristems were analyzed using confocal laser scanning microscopy. As expected, *in vivo* expression patterns highly correlated with reconstructed expression patterns of genes used as reference genes (*ETTIN; SHOOT MERISTEMLESS*, *STM* and *MERISTEM LAYER 1*, *ATML1*) as well as genes with high PEP scores, e.g. AT1G62500 (*CO2*, PEP = 0.17), while there was lower overlap with reconstructed expression patterns of genes with low PEP scores, such as *SHORT ROOT* (*SHR,* PEP = 0.15), and *PIN-FORMED 1* (*PIN1,* PEP = 0.14). In general, the prediction broadly recovered the cells and tissues that show activities of the genes, but some gene expression patterns were more restricted in the reporter gene analyses (e.g. *SHR*, *PIN1*). This could be explained by the limited set of reference genes that was used for the prediction, but also by the possibility that the reporter gene constructs do not contain all regulatory elements needed for correct spatial expression of the genes.

**Figure 3:**
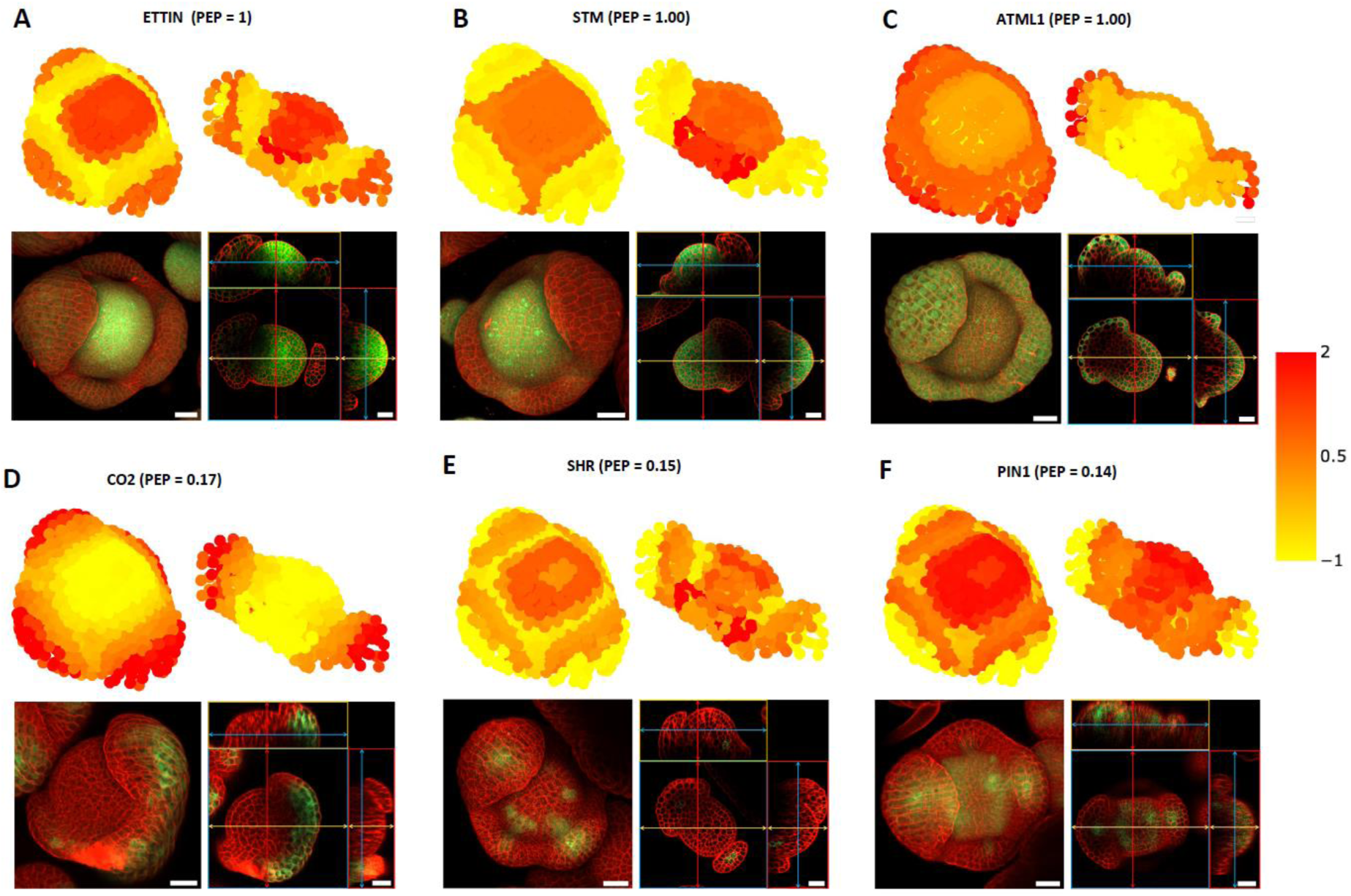
Validation of reconstructed gene expression patterns with reporter gene assays. Upper part in A-F shows the predicted expression of ETTIN, STM, ATML1, CO2, SHR and PIN1 from the top and cross section view of stage 4 flower meristems. Titles include gene symbol and PEP score for the predicted 3D expression profile. The lower part in A-F shows the GFP expression pattern (green) for plant lines under the control of the respective promoter, as detected by confocal laser scanning microscopy in A. thaliana stage 4 flower meristems. Confocal images show the flower meristem from the top (left) as well as different orthogonal sections (right). Cell walls were stained using propidium iodide (red). Scale bars indicate 20 µm.

### Gene expression reconstruction of floral meristem domains

Next, we evaluated the performance of spatial expression reconstruction to study quantitative gene expression in particular domains that give rise to the different organ types in the flower. In Arabidopsis flower development, the identities of different organ types are determined by floral homeotic transcription factors. In particular, sepals are specified by the activity of *APETALA1 (AP1)*, petals are defined by the combination of *AP1* and *APETALA3 (AP3)*, stamens are specified by *AP3* and *AGAMOUS (AG)*, and the carpels is determined by *AG* activity.

We estimated the expression of a gene in the *AP3-* and *AG-* domains of the 3D reconstructed meristem, as the average expression of that gene in the cells which had a positive expression of *AP3* or *AG* reference genes, respectively. To validate these results we generated sorted nuclei RNA-seq (FANS-RNA-seq) from floral meristems expressing nuclear targeting fusion protein (NTF) (Deal and Henikoff 2011) in *AP3* vs. *AG* expression domains. The GFP-containing NTF protein was transcribed under the control of *AP3* promoter (*pAP3*::NTF) and the second intron of *AG* (*pAGi*::NTF) in the floral induction system (*pAP1*:AP1-GR *ap1-1 cal-1*; Kaufmann et al. 2010). The expression patterns of *pAP3*::NTF and *pAGi*::NTF were visualized by confocal microscopy (Sup Fig 8), and the nuclei of *AP3*- or *AG*-expression domains were sorted based on the positive GFP signal in FANS.

Transcriptomes retrieved from the spatially reconstructed *AP3* and *AG* domains in the floral meristem showed a high correlation with the domain-specific bulk RNA-seq expression (*Rho*=0.89 for *AP3*- and *Rho*=0.88 for *AG*- domain when using genes with a PEP higher than 0.13). This was close to the correlation obtained among the bulk RNA-seq biological replicates (*Rho*=0.95 for *AP3* and Rho=0.93 for *AG*) when using the same set of genes (Fig. 4) which indicates a very good performance of the method. Even more interesting, the reconstructed expression was able to recover the log_2_ fold-change gene expression between both domains (Sup Fig 9A, Rho=0.37) when using genes with a PEP higher than 0.13 (n=1,306). In particular, the obtained correlation was very close to the correlation of the log_2_ fold-change gene expression obtained from the bulk RNA–seq biological replicates when using the same set of genes (*Rho*=0.47, Sup Fig 9C). This indicates that spatially reconstructed transcriptomes are able to predict domain-specific differential gene expression. The correlation between gene expression prediction and domain-specific bulk RNA-seq increases with increasing PEP scores (Sup Fig 9B), which is in agreement with the notion of the PEP score being an indicator of the quality for the predicted 3D expression.

**Figure 4.**
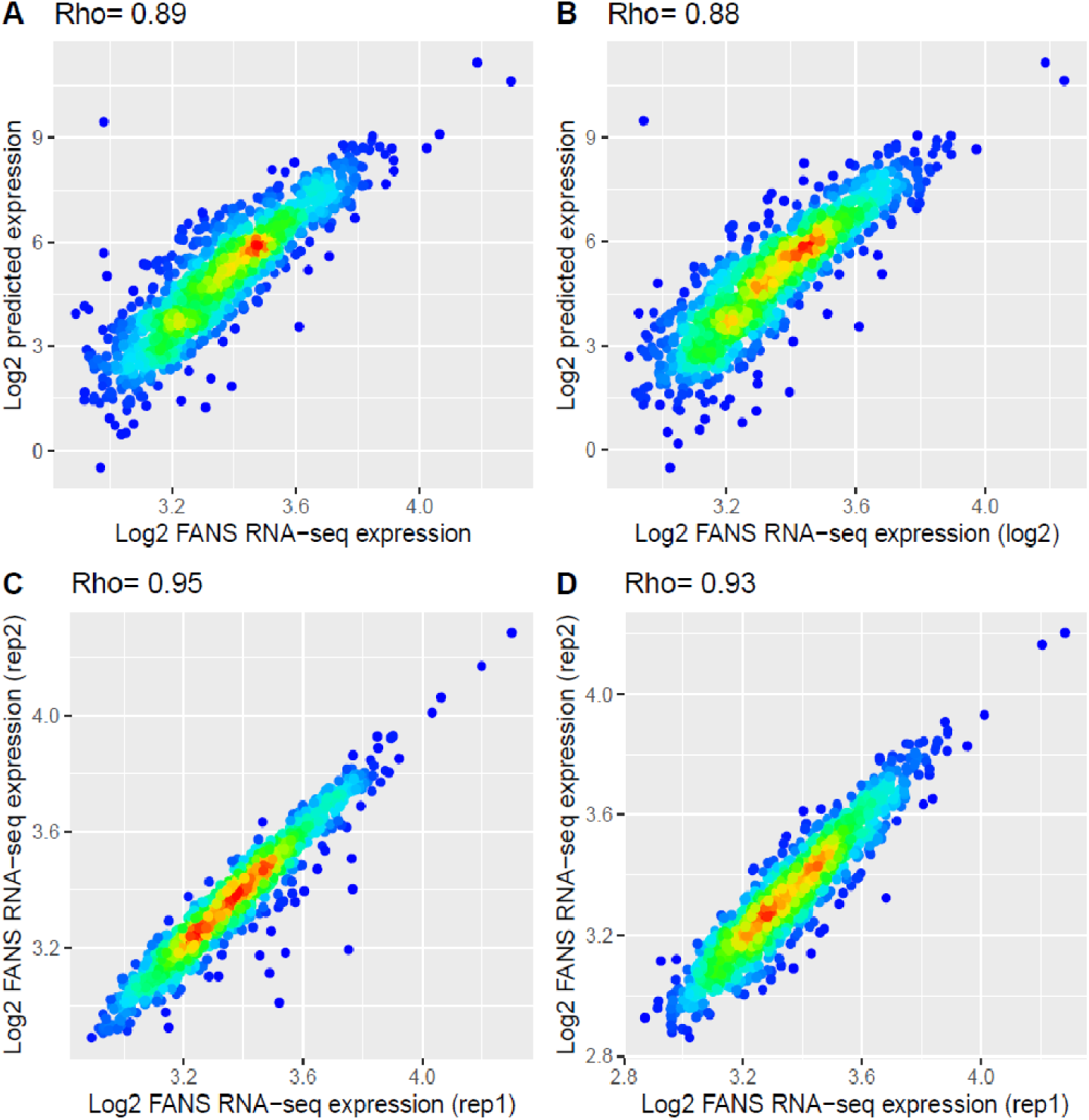
Prediction of AP3 and AG domain gene expression. Scatterplot showing the gene expression for AP3 (A) and AG (B) domain predicted by our method (y-axis) and observed by our FANS bulk RNA-seq data (x-axis) when using genes with PEP value>0.13 (n=1,306). Bottom row shows the scatterplot for the gene expression of both biological FANS bulk RNA-seq replicates for AP3 (C) and AG (D).

In this way, we detected a large number of genes with specific expression (Sup Table 2) in the different floral whorls as determined by the (combined) expression of *AP1* (sepal), *AP3-AP1* (petal), *AP3-AG* (stamen) and *AG* (carpel). For example, we predict a higher expression of *APETALA2* in the sepal domain, which is in line with its known role in sepal specification together with *AP1* (Kunst et al. 1989). We predict *PETAL LOSS (PTL)* expression in the *AP1* and *AP1*-*AP3* domain, which is consistent with previous findings that *PTL* is expressed in sepal margins while controlling petal development (Brewer et al. 2004). In the other hand, we predict *PERIANTHIA* to be strongly induced in the three inner whorls, as expected from the literature (Chuang et al. 1999), while we predicted *UNUSUAL FLORAL ORGANS* (*UFO*) to be expressed in the *AP3-AG* and *AP3-AP1* domain which fits with the observed expression in the petal and stamen whorls (Samach et al. 1999). This exemplifies the power of the method to identify whorl-specific genes. The predicted floral whorl-specific expression is significantly related to direct DNA-binding of flower domain-specific TFs in their regulatory regions (Sup Table 2, Sup Fig 10).

It is worth to note that we could apply a similar methodology directly to the snRNA-seq data (w/o 3D reconstruction), where average domain-specific expression is calculated as the average expression among the snRNA-seq transcriptomes of nuclei that have a positive expression of *AP3*, or *AG* for each domain respectively. However, the obtained fold-change expression has low agreement with the domain-specific bulk RNA-seq data (Rho=0.04 pv<0.14, Sup Fig 11) when using the same genes as before (PEP > 0.13). This indicates that the advantage of integrating the 4,395 transcriptomes of the snRNA-seq data into a physical map of 1,331 cells has the additional benefits of obtaining a more accurate estimate of gene expression per cell as it is calculated since it combines the information from several snRNA-seq transcriptomes.

In summary, the presented data demonstrates that our method can be used to create a genome-wide 3D gene expression atlas of a plant organ, and to correctly predict gene expression and gene fold change expression of particular morphological regions that was not possible with the snRNA-seq data alone.

### The origin of vascular cell identity in the floral meristem

Spatial reconstruction of transcriptomics data can be used to pinpoint the spatial location of cells characterized with a particular transcriptome signature (e.g. snRNA-seq cell clusters, ploidy levels (Bhosale et al. 2018), vascular cells (Shi et al. 2021)) by using an expression similarity-based method. For example, the initial establishment of vascular stem cell identity in the apical meristems is not well known (Sanchez, Nehlin, and Greb 2012). The transcriptomes of vascular tissues in inflorescence stems have been characterized by FANS bulk RNA-seq (Shi et al. 2021), including *SMXL5* (distal cambium) and *PXY* (proximal cambium). Therefore, assuming that the vascular tissues have similar transcriptomes in the inflorescence stem and in the floral meristems, we could predict the location of vascular stem cells on the reconstructed 3D meristem even when they cannot be distinguished anatomically. We indeed obtained a distinct distribution pattern of vascular stem cells (Sup Fig 11), where the cambium (*PXY* and *SMXL5*) localizes in the cell layers adjacent and just below the floral meristem with a radial disposition. Confocal imaging confirmed that *PXY* and *SMXL5* expression is initiated in cells just adjacent/below the apical meristem, but in a specific subset of cells (Sup Fig 12). This discrepancy could be due to the low number of reference genes used, which may not allow to have the needed resolution. Once these cells have been located, their transcriptome can be estimated as explained before, obtaining a good correlation (Rho=0.34-0.42; Sup Fig 13) when compared with the FANS bulk RNA-seq data. This information can be used in future work to characterize the molecular control and regulatory networks of initiation of vascular identity in the floral meristem.

NovoSpaRc outputs the probability of each snRNA-seq transcriptome as corresponding to a particular cell in the spatial map. Therefore, we can map the location of the identified snRNA-seq clusters (https://threed-flower-meristem.herokuapp.com) and visualize their physical location. In particular, cluster 1-(pro)cambium shows the same location adjacent/below the apical meristem as the one estimated by transcriptome similarity.

In summary, this shows the potential to integrate different features (e.g. cells differentiating to vascular tissues) into a common spatial map which can be used to associate with the spatial expression profiles.

## DISCUSSION

The identity and function of plant cells is strongly influenced by their precise location within the plant body (Xu et al. 2021). Therefore, to understand plant development at the molecular level, it is important not only to characterize the molecular status and dynamics of each individual cell but also to know their physical location in the plant. As stated in the introduction, spatial genomics in plants have been limited to profile only a limited number of genes per experiment. Here, we provide a proof of concept for a methodology to overcome this limitation by combining scRNA-seq/snRNA-seq with a 3D microscope-based reconstructed floral meristem. In this way, we are able to reconstruct the spatial expression of a large number of genes (>1,000) in their native spatial context. Moreover, we were able to quantitatively estimate the expression of these genes in particular morphological regions of the floral meristem. Future work should develop more dedicated statistical methods to test for gene expression differences on the 3D reconstructed structure. One possibility is to apply a re-sampling approach to the snRNA-seq data. We envision that by independently mapping multiple subsamples of the snRNA-seq data to the reference map, we will be able to estimate the variance of the gene expression which is needed to test for differential gene expression in different (groups of) cells.

The number of high-quality genes predicted is heavily dependent on the number and identity of genes present in the reference spatial map. Thus, we provide a PEP score that can be used to estimate the performance of the predicted expression for each gene, even before having generated the reference spatial map. In this way, this score can be used to select the minimum set of reference genes needed to obtain a good prediction of the spatial expression of a desired group of genes. Hence, this score helps in planning the design of a spatial genomics experiment whose data will be used as a spatial reference to predict the spatial expression of a set of genes.

This methodology has the potential to be applied to other types of -omics experiments. For example, we are already applying it to map scATAC-seq experiments into the 3D reconstructed floral meristem (data not shown). This offers the additional benefit to be able to integrate multiple single cells -omics data in their natural physical context. Indeed, an important problem is how to integrate multiple single cell -omics experiments (e.g. scRNA-seq and scATAC-seq data). The typical approach (Stuart et al. 2019) is to find anchors between genes and ATAC-seq regions that allow us to link the cells profiled independently in both types of experiments. We envision that independently mapping the scRNA-seq data and the scATAC-seq to a common spatial map will be an alternative way to integrate both types of experiments. In addition, we have shown that we can map transcriptional signatures of particular features (e.g. cells differentiating to vascular tissues) to the reconstructed spatial map, allowing to annotate or integrate additional experiments/data in the spatial map.

Furthermore, time-series scRNA-seq datasets could also be tackled with this approach. For example, when live imaging has been used to reconstruct the spatial map at different time points and cell segmentation and lineage tracking have been used to infer cell lineage in the spatial map (e.g. Refahi et al. 2021), the inferred cell lineage can be used to link the cells at different time points. Alternatively, when the plant structure at the different time points considered has similar morphology, the scRNA-seq data could be mapped to the spatial map of one particular time point. Otherwise, computational alignment of the spatial maps at each time-point will be required.

In summary, these results provide a primer for future initiatives to generate plant organ 3D atlases and for studies aiming to understand single cell -omics studies with respect to plant morphology and development.

## MATERIALS AND METHODS

### Plant material

*pAP1*:AP1-GR *ap1-*1 *cal-*1 plants were grown at 22 °C under long-day conditions (16 h light, 8 h dark) on soil. After plants bolted and reached the height of 2 cm to 5 cm, they were induced daily by applying the DEX-induction solution (2 μM Dexamethasone and 0.00016% Silwet L-77) to their main inflorescences. Around 20 inflorescences were collected and snap-frozen in liquid nitrogen on the fourth day after the first induction.

### Nuclei isolation

Inflorescences were gently crushed to pieces in liquid nitrogen using a mortar and a pestle and then transferred to a gentleMACS M tube. After liquid nitrogen evaporated totally, 5 ml of Honda buffer (2.5% Ficoll 400, 5% Dextran T40, 0.4 M sucrose, 10 mM MgCl_2_, 1 μM DTT, 0.5% Triton X- 100, 1 tablet/50 ml cOmplete Protease Inhibitor Cocktail, 0.4 U/μl RiboLock, 25 mM Tris-HCl, pH 7.4) was added to the tube. Nuclei were released at 4 °C by homogenizing the tissue on a gentleMACS Dissociator with a running program as described previously (Sunaga-Franze et al. 2021). The resulting homogenate was filtered through a 70 μm strainer, and another 5ml Honda buffer was applied onto the filter to collect the remaining nuclei. Nuclei were then pelleted by centrifugation at 1000 *g* for 6 min at 4 °C and then resuspended gently in 500 μl Honda buffer. The nuclei suspension was filtered again through a 30 μm strainer, diluted by adding 500 μl PBS buffer, and stained with 2 μM DAPI. Ambion RNase Inhibitor and SUPERaseIn RNase Inhibitor were added to a final concentration of 0.4 U/μl and 0.2 U/μl, respectively. 200,000 events of single nuclei were selected on DAPI signals by a BD FACS Aria Fusion into a 1.5-ml tube with landing buffer (15 μl 4% BSA in PBS with 80 U Ambion RNase Inhibitor and 80 U SUPERaseIn RNase Inhibitor). Sorted nuclei were counted in Neubauer counting chambers under a Leica DMi8 fluorescent microscope.

### Preparation of snRNA-seq libraries

Single-nuclei RNA-seq library was prepared from 10,000 freshly-isolated plant nuclei with the Chromium Single Cell 3ʹ Reagent Kits v3 according to the manufacturer’s instructions. 14 PCR cycles were used for cDNA amplification, and 13 PCR cycles were used for final amplification of the constructed libraries. The average fragment size of the snRNA-seq library was checked with an Agilent High Sensitivity D1000 ScreenTape, and the concentration was measured with Qubit 1X dsDNA HS Assay Kit. Sequencing was performed on a HiSeq 4000 (Illumina) platform.

### Preparation of domain-specific RNA-seq libraries

Nuclei were from *pAP1*:AP1-GR *ap1*-1 *cal*-1 transgenic plants expressing a GFP labeled nuclei envelope protein driven by tissue-specific promoters (Deal and Henikoff 2011). We used *AP3* promoter and *AG* 2nd intron plus a minimal 35S promoter element as promoters for the constructs to mark *AP3* and *AG* expressing domains in flowers, respectively. After nuclei isolation, as described in the previous paragraph, nuclei were sorted into a 1.5-ml tube with 15 μl of 4% BSA in PBS and 6 μl of RiboLock RNase Inhibitor by a BD FACS Aria III. The GFP channel was set using *pAP1*:AP1-GR *ap1-*1 *cal-*1 as a negative control, and then nuclei were selected by gating on the DAPI peaks under the GFP positive events. After sorting, nuclei were pelleted at 1500 *g* for 10 min at 4 °C, and the supernatant was then removed. Nuclei were lysed by vortex in 350 μl RLT buffer with 2-Mercaptoethanol, and RNA was then isolated with Qiagen RNeasy Micro Kit. After RNA isolation, cDNA synthesis was done with SMART-Seq® v4 Ultra® Low Input RNA Kit following the manufacturer’s instructions. cDNA was sheared to 200–500 bp size by Covaris AFA system and constructed with sequencing adaptors by ThruPLEX DNA-Seq Kit.

### Confocal imaging

GFP expressing plant lines under the control of the *CO2* (AT1G62500), *PIN1* (AT1G73590) and *SHR* (AT4G37650) promoters were obtained from the Nottingham *Arabidopsis* Stock Centre (NASC, UK) as part of the SWELL line seed collection (BREAK line set N2106365), which was previously described in roots by Marquès-Bueno *et al*. (2016). To generate plant lines driving GFP expression from the *ETT/ARF3* (AT2G33860) promoter, we inserted a 3 kb long promoter fragment into the pK7GW-INTACT_AT vector (Ron *et al*., 2014) using gateway cloning. Similarly, the 6.1 kb promoter of *STM* (AT1G62360) and the 5 kb promoter of *ATML1* (AT4G21750) were introduced into the pK7GW-INTACT_AT vector. *A. thaliana* Col-0 wild type plants were transformed by floral dip method (Clough and Bent, 1998). Plant lines expressing HISTONE 4 (H4)-coupled GFP under the control of the *PXY* (AT5G61480) and the *SMXL5* (AT5G57130) promoters (PXY:H4-GFP/SMXL5:H4-GFP) were previously described in Shi et al. 2021. For GFP expression analysis, plants were grown on soil at 22 °C and 16/8 h light/dark cycles using daylight led lights (200 μmol·m−2·s−1).

GFP expression was detected by confocal laser scanning microscopy using the Zeiss LSM 800 confocal microscope equipped with a Plan-Apochromat 20x/0.8 M27 or a C-Apochromat 40x/1.2 W Korr objective. GFP was excited at a wavelength of 488 nm with an argon laser, while emission was filtered by a 410-532 nm band pass filter. Propidium iodide (Sigma-Aldrich) was used to stain cell walls. It was excited at a wavelength of 305 nm and detected in a range of 595-617 nm. Z-stack images were median corrected and merged to orthogonal projections using the ZEN imaging software (Zeiss).

### snRNA-seq data analysis

Fastq files were processed with CellRanger v3.1.0 with default parameter values and using the Araport11 gene annotation (Cheng et al. 2017), obtaining 7,716 nuclei transcriptomes as a read count matrix. Genes encoded in the organelles were removed. Next, read count normalization and clustering were done with the R package Seurat v3.2.3 (Stuart et al. 2019). In particular, nuclei transcriptomes with less than 1,000 expressed were removed and SCT-normalization was applied within the SEURAT package setting the parameter *variable.features.n* to 3,000 and other parameters to default values. Next, the optimal number of PCAs was chosen to be the first 9 principal components by plotting the standard deviations of the principal components using the *RunPCA* and *ElbowPlot* functions. UMAP dimensionality reduction was obtained with the *runUMAP* function using the parameters *values dims = 1:9, reduction = ’pca’, n.neighbors = 50, min.dist = 0.01, umap.method = “uwot”, metric = “cosine”*. In order to identify clusters in the UMAP space, we used *FindNeighbors* and *FindClusters* functions with parameter values *resolution* = 0.04, *algorithm* = 1 and default values for other parameters. Marker genes for each cluster were identified with the function *FindAllMarkers* and parameter values: *only.pos = TRUE, assay=“SCT”, slot=“scale.data”, min.pct = 0.25, logfc.threshold = 0.25*. In order to annotate the identified clusters, the average relative expression of the top 20 cluster marker genes in different publically available RNA-seq (see bulk RNA-seq analysis) and microarray samples were visualized in heatmaps in order to help to annotate the clusters. Expression values for GSE28109 (Yadav et al. 2014) were downloaded directly from GEO omnibus (file: GSE28109_averaged_mas5_data.txt). Heatmaps showing the expression of markers genes were calculated as the average relative expression across all nuclei for each cluster. Relative expression was calculated as the normalized read count expression of a gene minus the average expression of this gene across all samples/nuclei considered.

### Bulk RNA-seq analysis

Fastq files from publicly available bulk RNA-seq data were downloaded from Sequence Read Archive (SRA; https://www.ncbi.nlm.nih.gov/sra). The next analysis was done for each dataset independently. The analyzed datasets were: PRJNA314076 (Klepikova et al. 2016), PRJNA471232 (Tian et al. 2019); PRJNA595605 (Shi et al. 2021), and the *AG*- and *AP3*- domain specific bulk RNA seq data generated in this project. Fastq files were trimmed from adapters using Trimmomatic v0.36 (Bolger, Lohse, and Usadel 2014). The reads were then mapped to the TAIR10 Arabidopsis genome using STAR v2.7.0b (Dobin et al. 2013) with parameter values -*-alignIntronMax 10000 -- outFilterMultimapNmax 1 --outSJfilterReads Unique* and other parameters with default values. *featureCounts* (Liao, Smyth, and Shi 2014) was used to count the number of mapped reads per gene (in exon and introns) with default parameters. Next, reads mapping to genes encoded in the organelles were removed. Only genes with more than 10 reads mapped in at least 2 samples were considered in the further analyses. Read count data was analyzed with DESeq2 v1.24.0 (Love, Huber, and Anders 2014), in particular normalized expression was calculated with *varianceStabilizingTransformation* function using default parameters.

### snRNA-seq and spatial gene expression map data integration

snRNA-seq data was processed as described in the previous section, which results in a matrix of normalized expression values of 6,104 nuclei and 19,718 genes, genes expressed in less than 30 cells were removed (n=2,890) with the exception of *WUS* and *CLV3* which were kept in the dataset due their biological importance. Data of the spatial map containing positional coordinates of 1,451 cells, their associated cell growth, cell volume, lineage and expression of 28 genes for the reconstructed 3D stage 4 floral meristem was downloaded from Refahi et al. 2021. First, cells (n=52) with expression of none of the 28 reference genes were removed. Next, genes (n=5) with the same expression in all nuclei or not present in the normalized snRNA-seq dataset were removed as they are not informative for the data integration procedure. Cells from the spatial map (n=68) were removed when they had less than 3 reference genes expressed, or when the combination of genes expressed in one particular cell was present in less than 4 other cells. This resulted in a spatial map of 1,331 cells and 23 genes. Next, nuclei from the snRNA-seq datasets not expressing any of the 23 genes considered in the reconstructed meristem were removed. At this step, the snRNA-seq contained 5,910 nuclei and 16,828 genes. The resulting snRNA-seq dataset and the reconstructed floral meristem were integrated using NovoSpaRc v0.4.1 (Nitzan et al. 2019). As described in the main text, 3 modifications were considered:

1. **Filtering.** When this modification was applied, distances between all the transcriptomes of the snRNA-seq and the spatial map were calculated. Only the top 50 snRNA-seq transcriptomes with closest distance to each cell of the spatial map were kept in order to eliminate nuclei that were not present in the spatial map (e.g. cauline leaves, pedicel…). The final number of snRNA-seq nuclei depends on the distance used.
2. **Genes used for calculating distance among the snRNA-seq transcriptomes.** The standard NovoSpaRc procedure uses the highly variable genes identified by the program to analyze the snRNA-seq data in order to calculate the distances among the snRNA-seq transcriptomes. We modified this option to use the top 100 genes with the highest pearson correlation value in the snRNA-seq space to the 23 genes considered in the spatial map. In our case, this results in 1,709 unique genes.
3. **Distance**. By default, NovoSpaRc used the Euclidean distance between the snRNA-seq and spatial map transcriptomes. We also included Jaccard and Hamming distances for binary data. When these distances were used, the snRNA-seq data was binarized as non-expressed when the normalized expression of a gene was zero and as expressed when the normalized expression was bigger than zero. When using the Euclidean distance, we include the optional binarization of the snRNA-seq expression data.

The best set of modifications and parameter value sets was chosen as the ones minimizing the average AUCROC of the genes from the spatial map except *AHP6*, *ETT*, *WUS* and *CLV3*, we excluded these 4 genes because their performance was always poor independently of the parameter values used and/or because the low number of cells where they were expressed in the spatial map. The final parameter set was using all three proposed modifications, in particular using the Jaccard distance, and with values for the NovoSpaRc parameters: *num_neighbors_source=2, num_neighbors_target=5, epsilon = 0.05, alpha =.1, max_iter=5000* and *tol=1e-9.* As output, NovoSpaRc provides a matrix (Gromoth-Wasserstein matrix, GW) containing the probabilistic assignment of each nucleus from the snRNA-seq to each of the cells of the spatial map. For numerical reasons (to avoid long decimals), the GW matrix was multiplied by 10^5^. It also outputs the predicted expression of each gene considered in the spatial map space.

### PEP score calculation

The Spearman correlation coefficient for a particular gene against each reference gene was calculated in the scRNA-seq data after the filtering step. The highest Spearman correlation coefficient was chosen as the PEP score for that particular gene.

### Localization of the vascular stem cells into the spatial map

FANS RNA-seq data (Shi et al. 2021) was analyzed as explained above. After, the data was log2 transformed, and the expression of each gene was normalized to have mean 0. The same procedure was applied to the gene expression profiles of the spatial map. Pearson correlation was calculated between each FANS RNA-seq dataset to transcriptome of each cell of the spatial map. Only genes (n=1,281) defined as vascular markers in (Shi et al. 2021) were used to calculate the correlation. P-values were calculated by testing if the correlation was higher than zero.

### Localization of the snRNA-seq clusters into the spatial map

NovoSpaRc outputs the probability of each snRNA-seq transcriptome as corresponding to a particular cell in the spatial map (GW matrix). Once obtained, the GW matrix was transformed so that columns (corresponding to cells in the spatial map) sum to 1. The score of one cell of the spatial map belonging to a particular cluster was calculated as the sum of the probabilities of all snRNA-seq transcriptomes of one particular cluster belonging to that particular cell in the spatial map.

## Supporting information

Supplementary Table 1. List of marker genes and annotation of the snRNA-seq clusters

Supplementary Table 2. Gene expression prediction in the floral whorls.

## DATA STATEMENT

The snRNA-seq and bulk RNA-seq data have been deposited in the GEO database under accession number GSE174599 and GSE174656, respectively.

## FUNDING

The work was supported by DFG (grant no. KA 2720/5-1 to X.X, K.K, and grant KA 2720/9-1 to M.M. K.K.), by an ERC Consolidator grant (PLANTSTEMS, #647148) to T.G.

## CONFLICT OF INTEREST

The authors declare no competing interests.

## ACKNOWLEDGEMENTS

We thank Johanna Müschner for their technical support. This work was supported by the BMBF-funded de.NBI Cloud within the German Network for Bioinformatics Infrastructure (de.NBI).

## SUPPORTING INFORMATION

### Supplementary Tables

Supplementary Table 1. List of marker genes and annotation of the snRNA-seq clusters.

Supplementary Table 2. Gene expression prediction in the floral whorls.

## SUPPLEMENTARY FIGURES AND LEGENDS

**Sup Figure 1:**
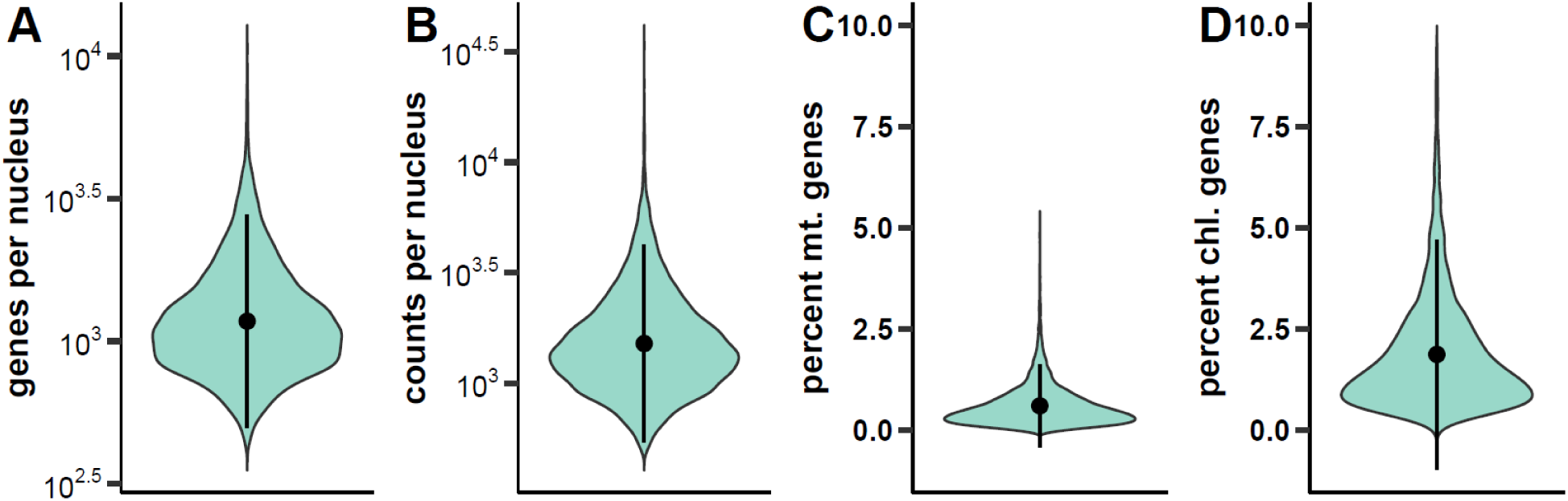
Summary of main statistics of the snRNA-seq. A) Number of expressed genes (containing at least one read) per nucleus. B) Number of mapped reads per nucleus. C) Percentage of reads mapping to the mitochondrial genome. D) Percentage of reads mapping to the chloroplast genome.

**Sup Figure 2.**
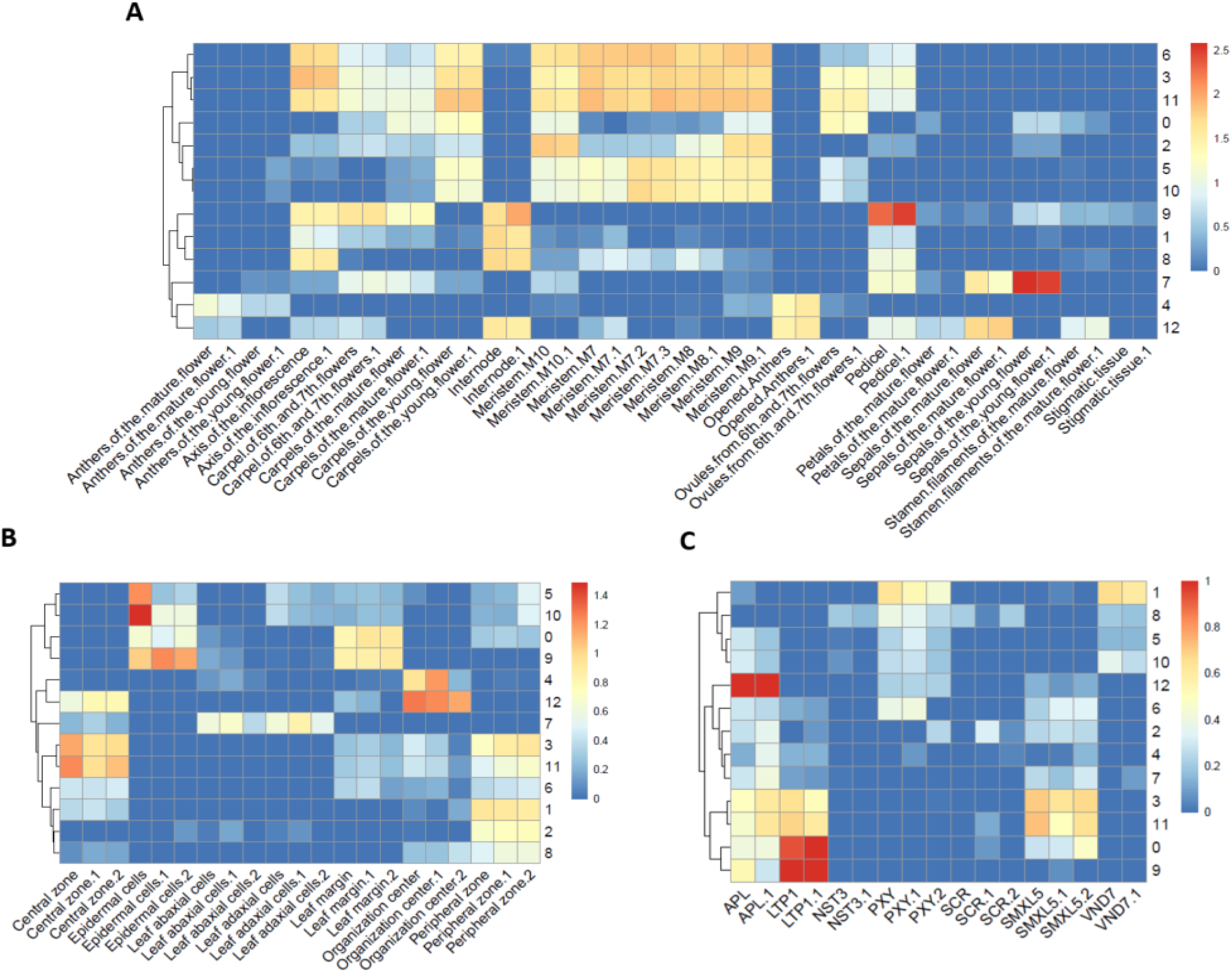
Average expression of the top 20 marker genes in publically available bulk RNA-seq datasets. Heatmaps show the expression of the top 20 significant marker genes for each snRNA-seq cluster in different publically available bulk expression profiles of: several flower organs and developmental stages (Klepikova et al. 2016) (A), shoot apical meristem domains (Tian et al. 2019) (B), and vascular tissues of inflorescence stems (Shi et al. 2021) (C).

**Sup Fig 3.**
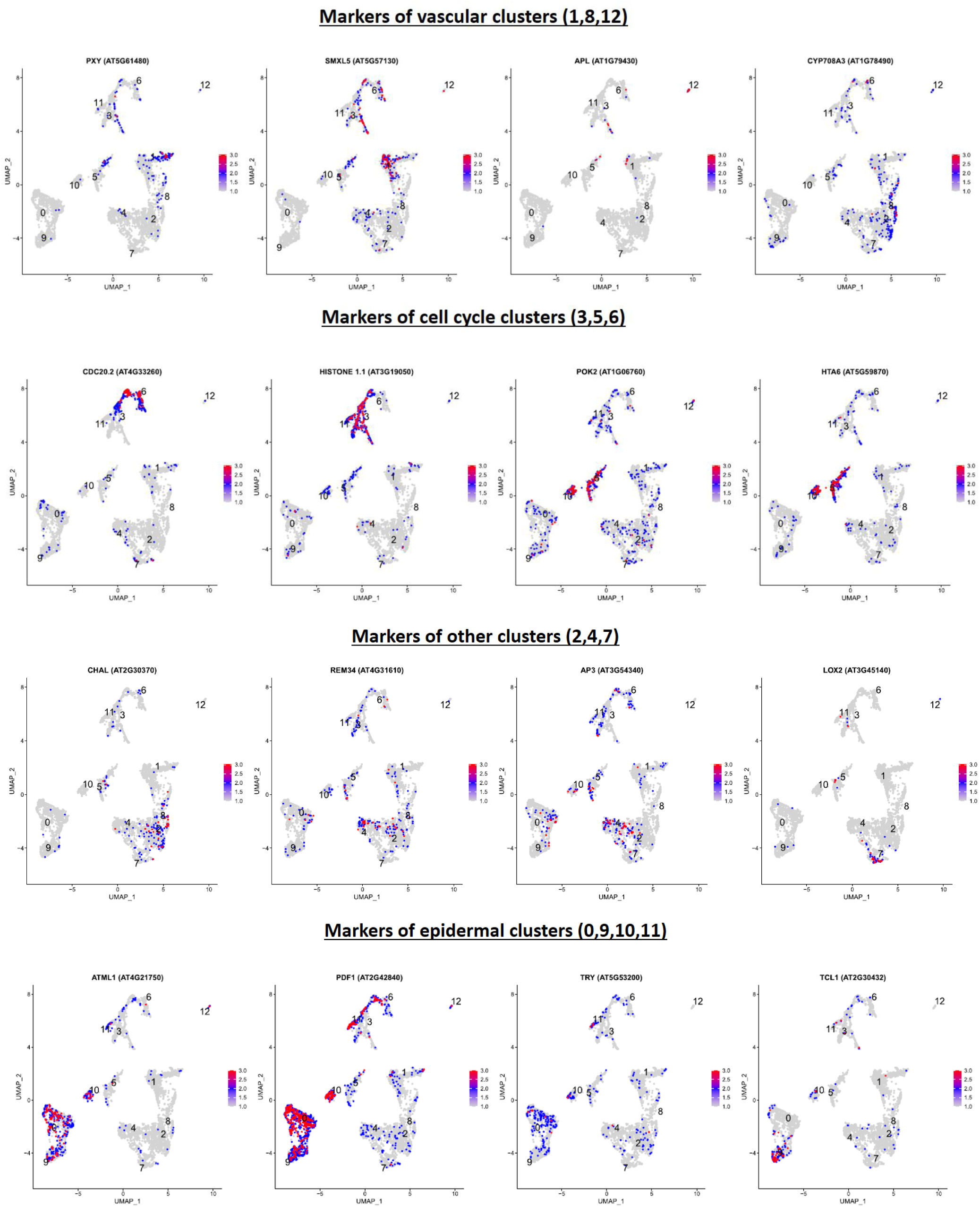
Expression of selected marker genes of snRNA-seq clusters on the UMAP plot.

**Sup Fig 4.**
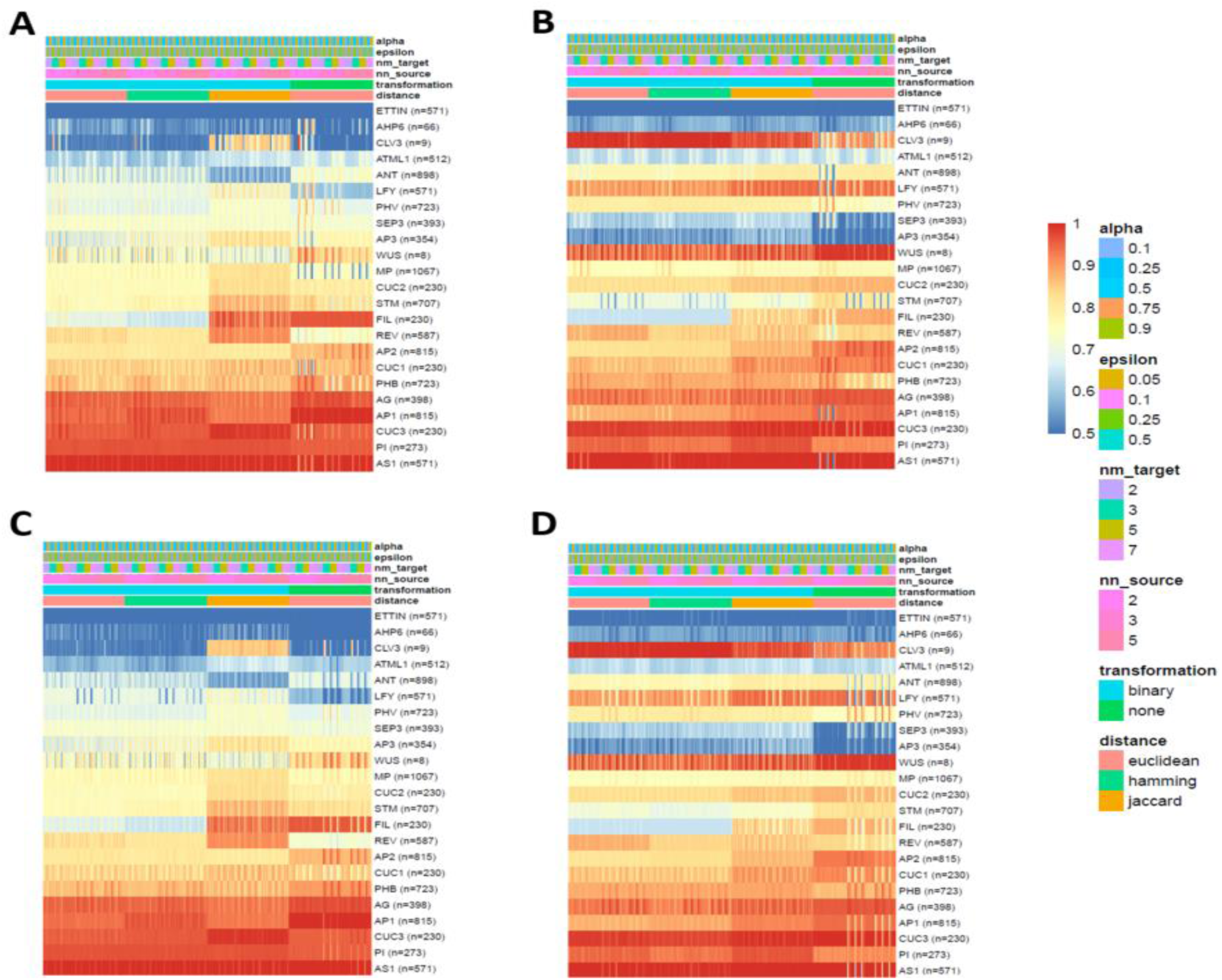
Gene-based performance of the method for gene expression reconstruction. Heatmaps show the performance (AUROC) for each reference gene when that particular gene was removed from the spatial map during the data integration step. Four models were tested: A) Filtering out snRNA-seq nuclei too dissimilar to the spatial map in the transcriptomic space (see Material and Methods) and using genes with high correlation with the reference genes in order to calculate transcriptomic distance among snRNA-seq nuclei (see Material and Methods). B) Applying no filter to the snRNA-seq and using genes with high correlation with the reference genes in order to calculate transcriptomic distance among snRNA-seq nuclei. C) Filtering out snRNA-seq nuclei too dissimilar to the spatial map in the transcriptomic space, and using the set of high variable genes defined by SEURAT to calculate transcriptomic distances between snRNA-seq. D) Applying no filter to the snRNA-seq data and using the set of high variable genes defined by SEURAT to calculate transcriptomic distances between snRNA-seq, this is the original option in novoSpaRc. The number between parentheses after the gene symbol indicates the number of cells where the particular gene is expressed in the spatial map. Legend indicates the different parameter values used for running novoSpaRc.

**Sup Fig 5.**
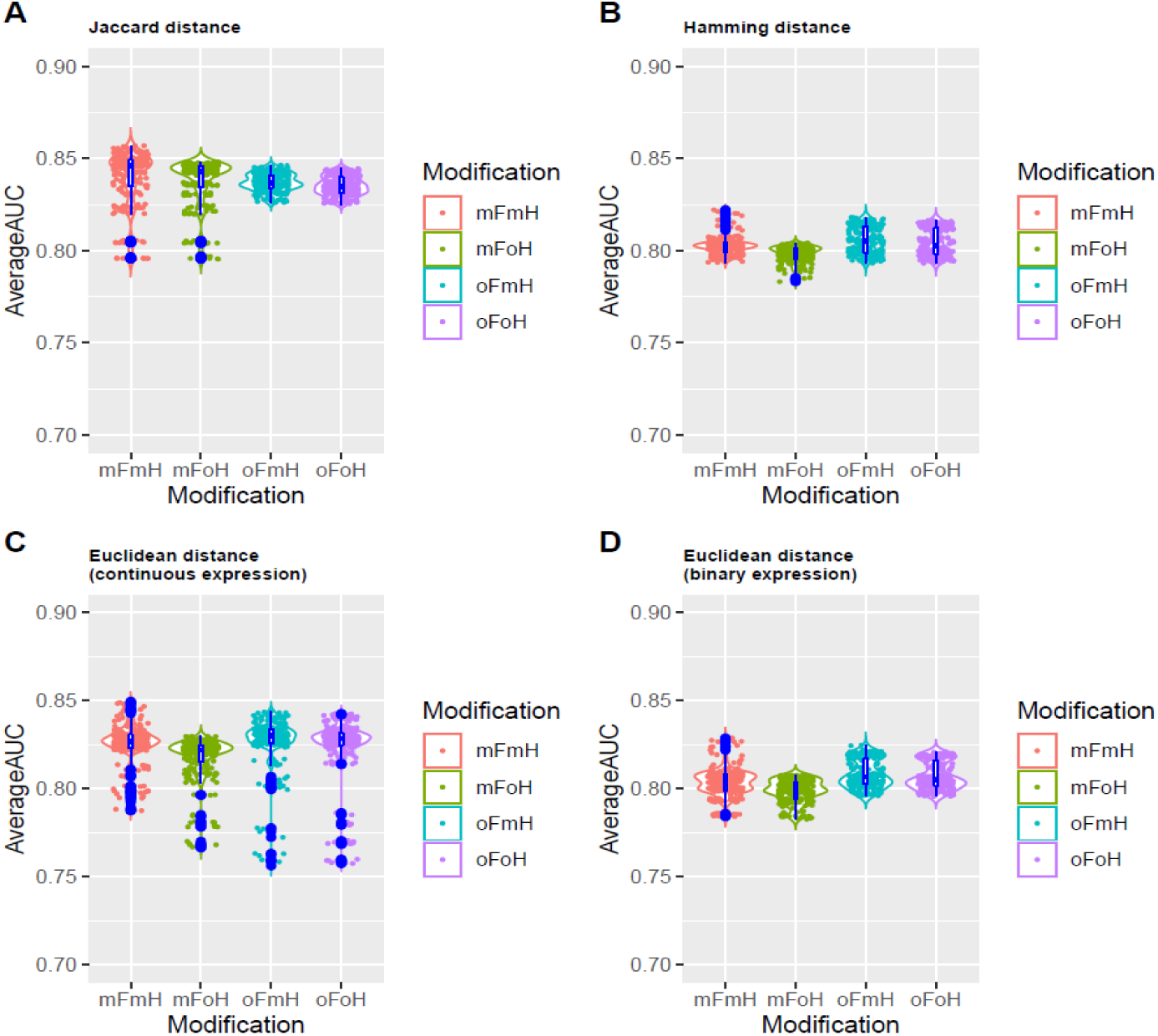
Average performance of our method for gene expression reconstruction. Violin plots show the average AUROC values across all reference genes except ETTIN, AHP6, CLV3 and WUS, which were excluded because of their consistent low performance or because of the low number of cells where they are expressed. Four distances were tested: Jaccard (A), Hamming (B), Euclidean using snRNA-seq continuous expression (C) and Euclidean when the snRNA-seq data was binarized. For each distance, four models were tested: 1) Filtering out snRNA-seq nuclei too dissimilar to the spatial map in the transcriptomic space (see Material and Methods) and using genes with high correlation with the reference genes in order to calculate transcriptomic distance among snRNA-seq nuclei (mFmH). 2) Applying no filter to the snRNA-seq and using genes with high correlation with the reference genes in order to calculate transcriptomic distance among snRNA-seq nuclei (oFmH). 3) Filtering out snRNA-seq nuclei too dissimilar to the spatial map in the transcriptomic space, and using the set of high variable genes defined by SEURAT to calculate transcriptomic distances between snRNA-seq (mFoH). 4) Applying no filter to the snRNA-seq data and using the set of high variable genes defined by SEURAT to calculate transcriptomic distances between snRNA-seq, this is the original option in novoSpaRc (oFoH).

**Sup Fig 6.**
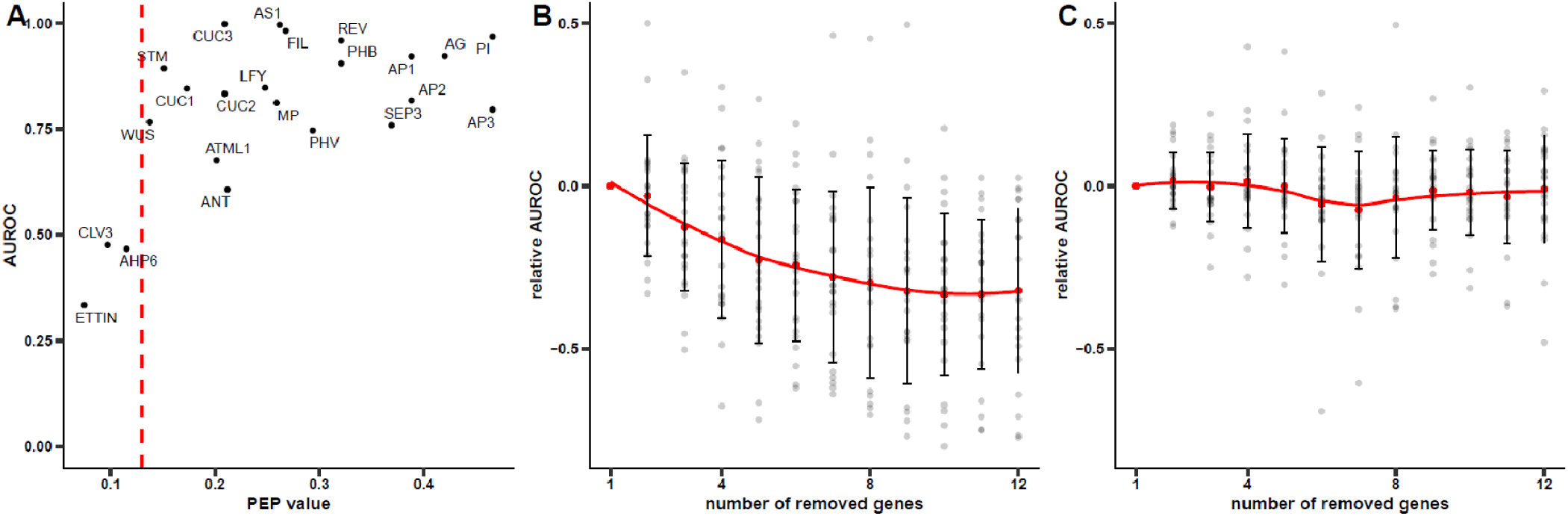
Performance of the reconstructed expression depends on PEP. A) Relationship between PEP and the performance (AUROC) of the gene expression estimation when the estimated gene was not included in the spatial map. Red line indicates the value 0.13. B) Performance (AUROC) of the prediction for each gene (grey points) when x genes from the spatial map with highest co-expression values with the predicted gene were sequentially removed. C) Performance (AUROC) of the prediction for each gene (grey points) when x genes from the spatial map with lowest co-expression values with the gene with the gene evaluated are removed. In B and C, the number of genes (x) removed is shown in the x-axis, and the drop in AUROC is shown in the y-axis; the red line represents a smoothing function (LOESS) applied to the average relative AUROC. Error bars indicate standard deviation.

**Sup Fig. 7:**
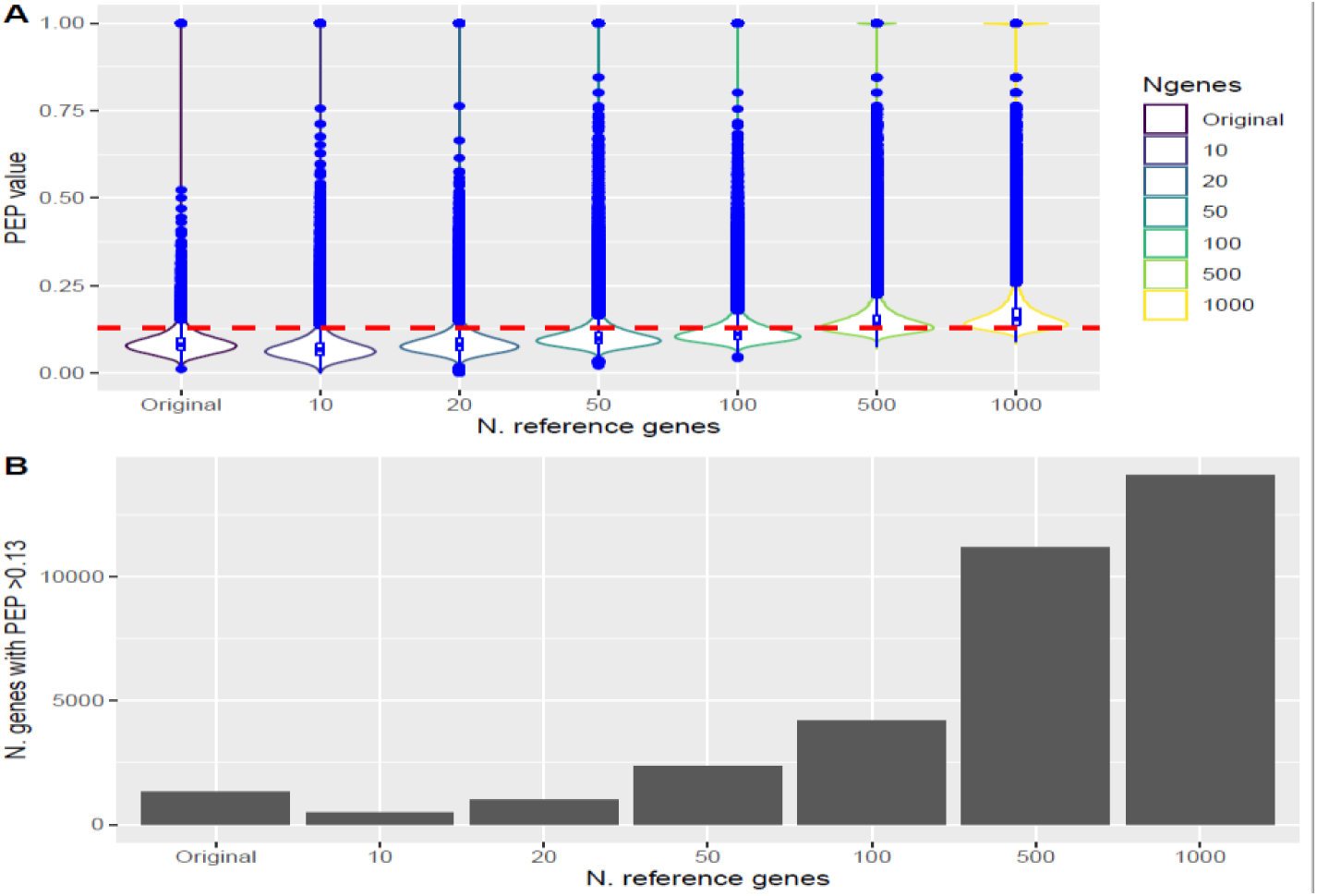
Increasing the number of reference genes increases the number of genes with PEP>0.13. A) PEP score distributions when using a random set on n reference genes (x-axis). B) Average number of genes with a PEP-score>0.13 depending of the number of reference genes used (x-axis).

**Sup Fig 8.**
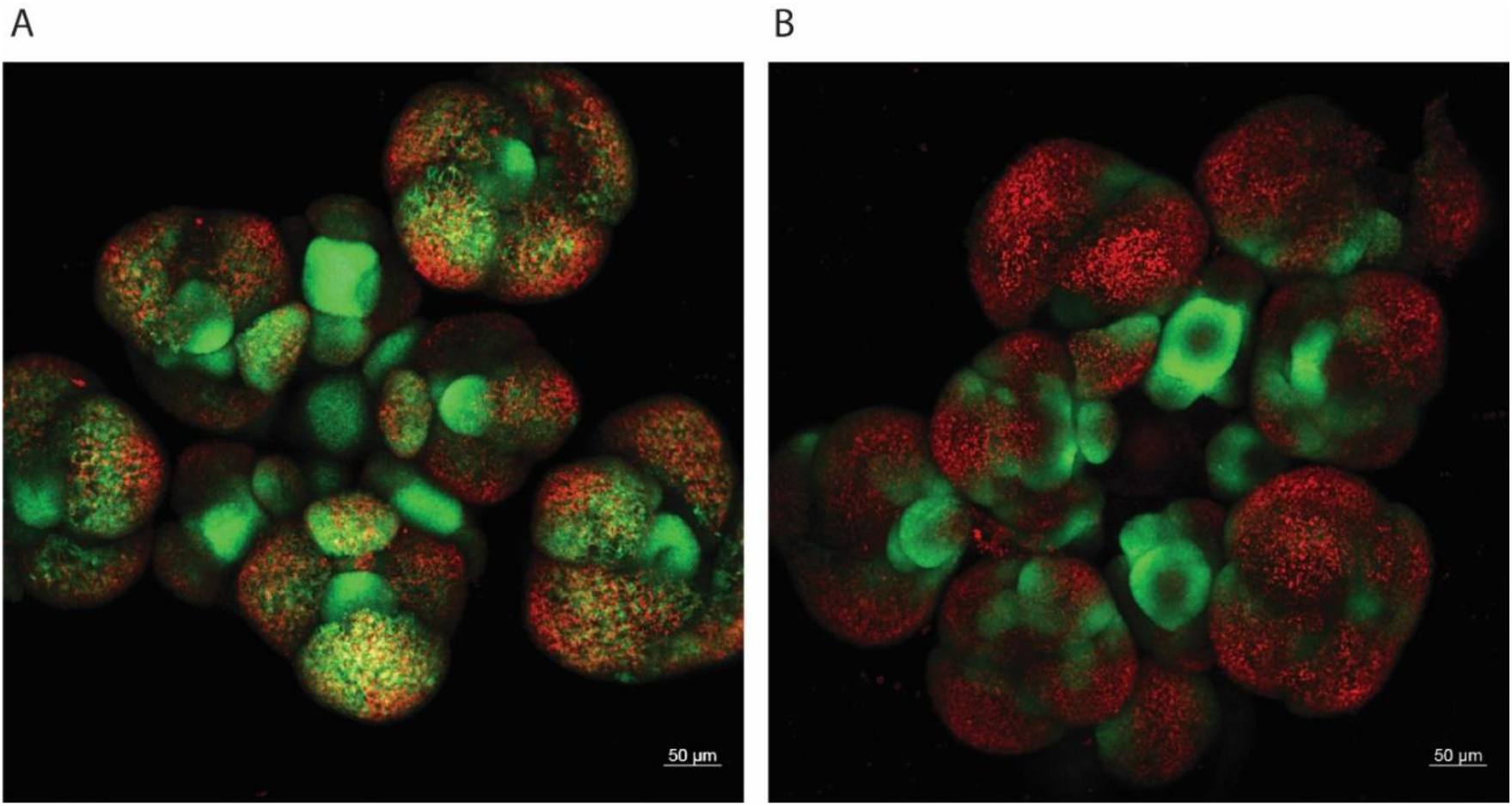
GFP signals in pAGi::NTF (A) and pAP3::NTF(B) domain specific lines used for FANS. Scale bars indicate 50 μm.

**Sup Fig 9.**
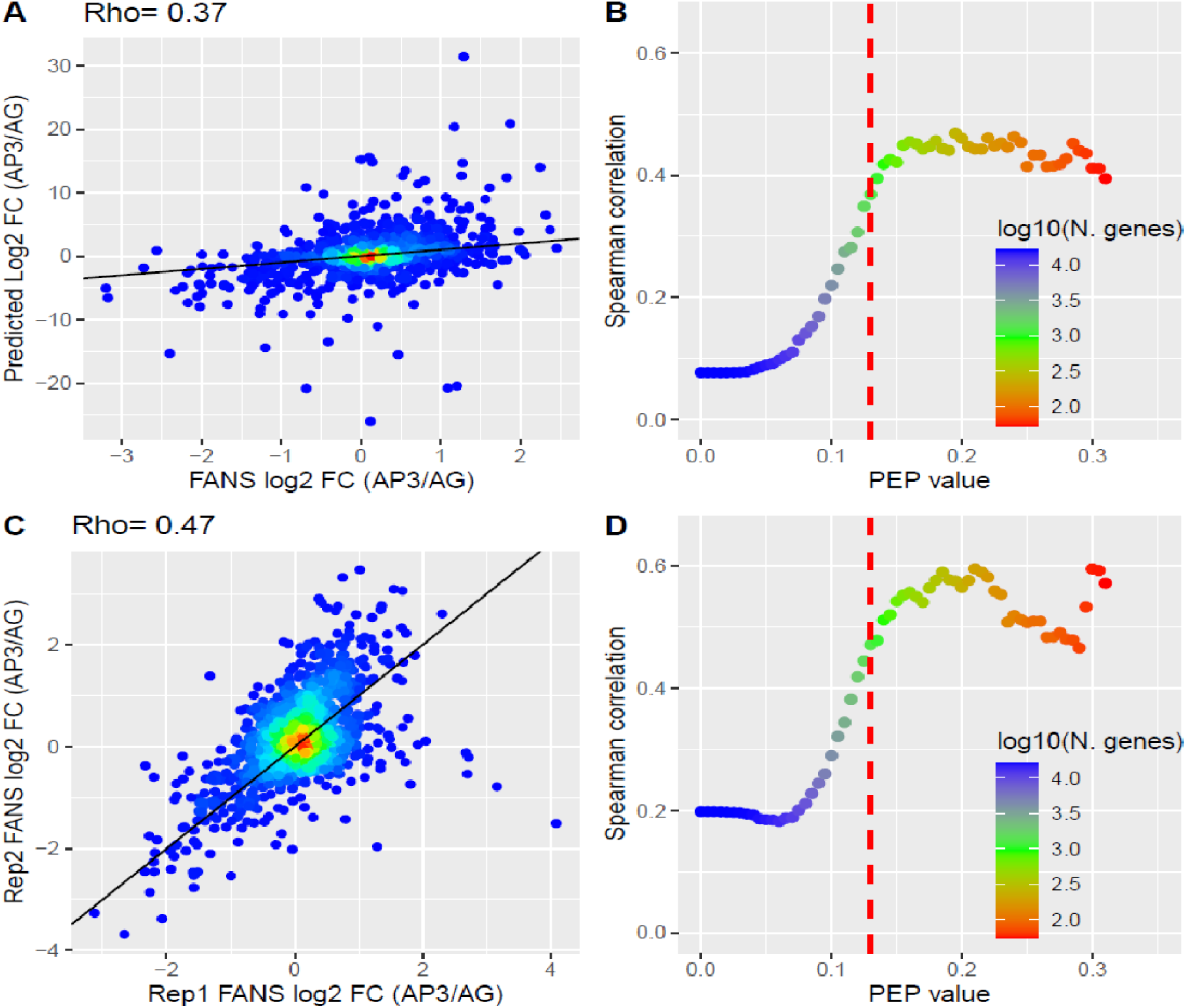
Prediction of AP3 vs AG domain-specific log2FC expression. A) Scatterplot showing the predicted change in expression between the AP3 and AG domain predicted by our method (y-axis) and observed by our bulk FANS RNA-seq data (x-axis) when using genes with PEP>0.13 (n=1,306). Continuous black line indicates the diagonal line. The associated Spearman correlation is 0.37. The associated spearman correlation for other values of PEP can be seen in B). Bottom row shows the scatterplot for the observed change in expression of both biological FANS bulk RNA-seq replicates for AP3 versus AG. Color in B and C indicates the number of genes predicted at this level of PEP. Vertical red line indicates the value of 0.13.

**Sup Fig 10.**
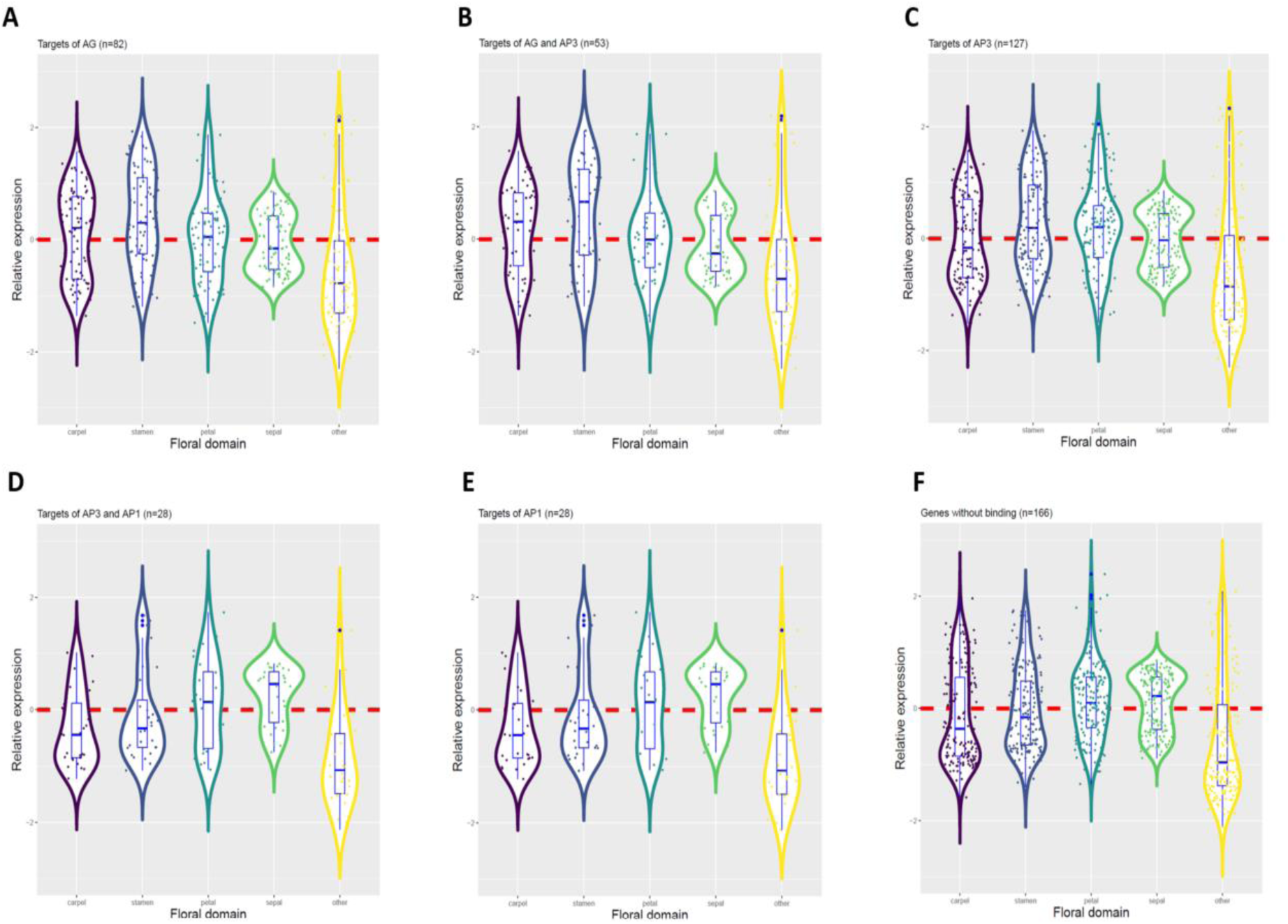
Gene expression distribution in the floral meristem whorls depending on TF binding. Gene expression was standardized to mean 0 variance 1, after average expression was calculated for each gene in the different floral whorls. Floral whorls are defined as: carpel: cells expressing AG but not AP3 neither AP1; stamen: cells expressing AG and AP3 but not AP1, petals: cells expressing AP3 and AP1 but not AG; sepals: cells expressing AP1 but not AG neither AP3. Four groups of genes were considered: A) genes with a AG binding in the gene body or the 2 kb regions around, B) genes with AG and AP3 binding, C) genes with an AP3 binding, D) genes with an AP3 and AP1 binding, E) genes with an AP1 binding and F) genes without any AG, AP3, or AP1 binding. Note that binding events of several TFs to the same gene does not necessitate that these TFs bind as part of the same complex, their binding could be independent and occur in different cells.

**Sup. Figure 11.**
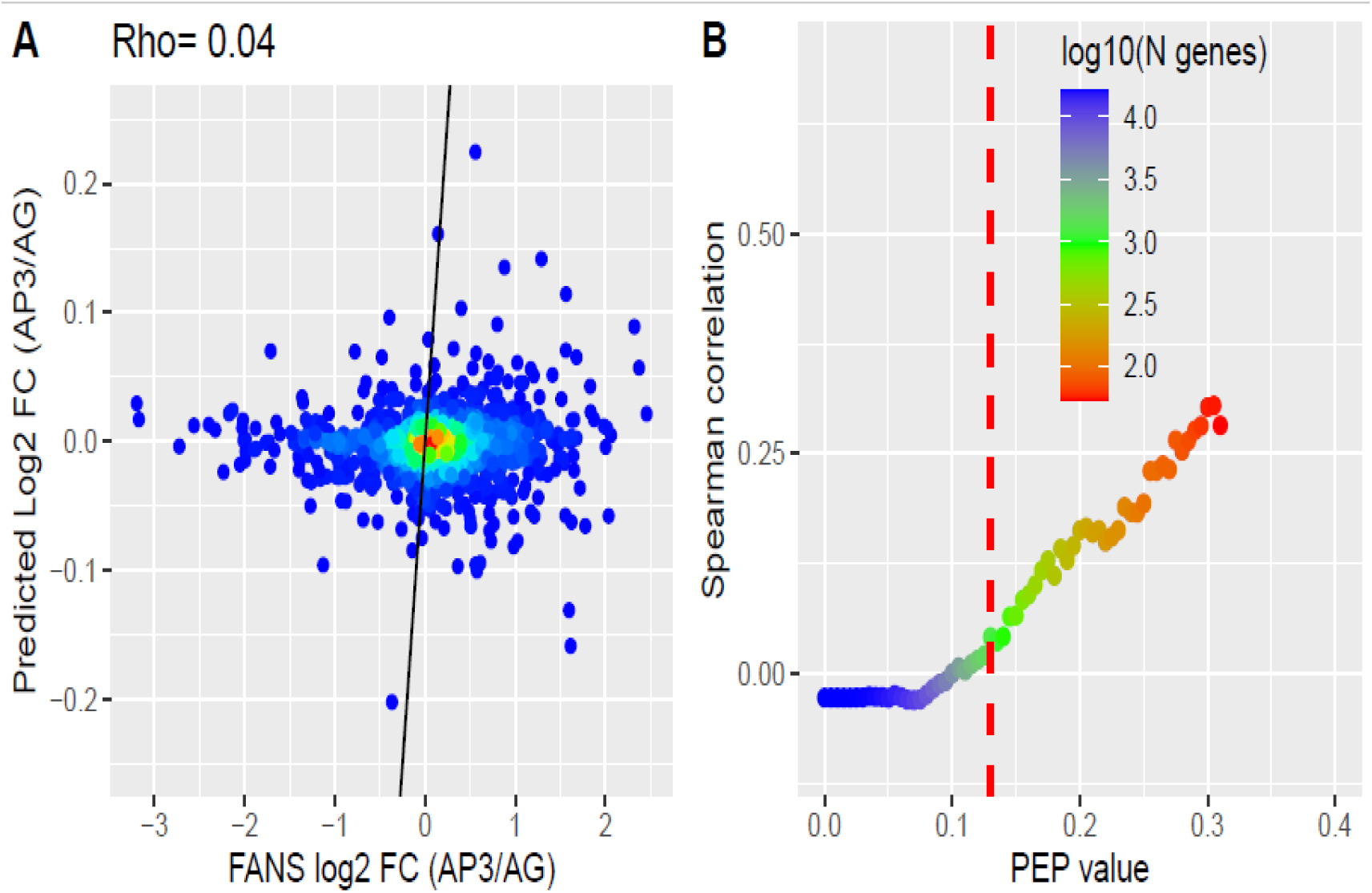
Prediction of AP3 vs AG domain-specific log2 FC expression directly from snRNA-seq. A) Scatterplot showing the predicted change in expression between the AP3 and AG domains predicted directly by the snRNA-seq (y-axis) or observed by our bulk RNA-seq data (x-axis) when using genes with PEP>0.13 (n=1,306). The associated Spearman correlation is 0.04 (pv< 0.14; not significant). The associated spearman correlation for other values of PEP can be seen in B). Vertical red line indicates the value of 0.13.

**Sup Fig 12.**
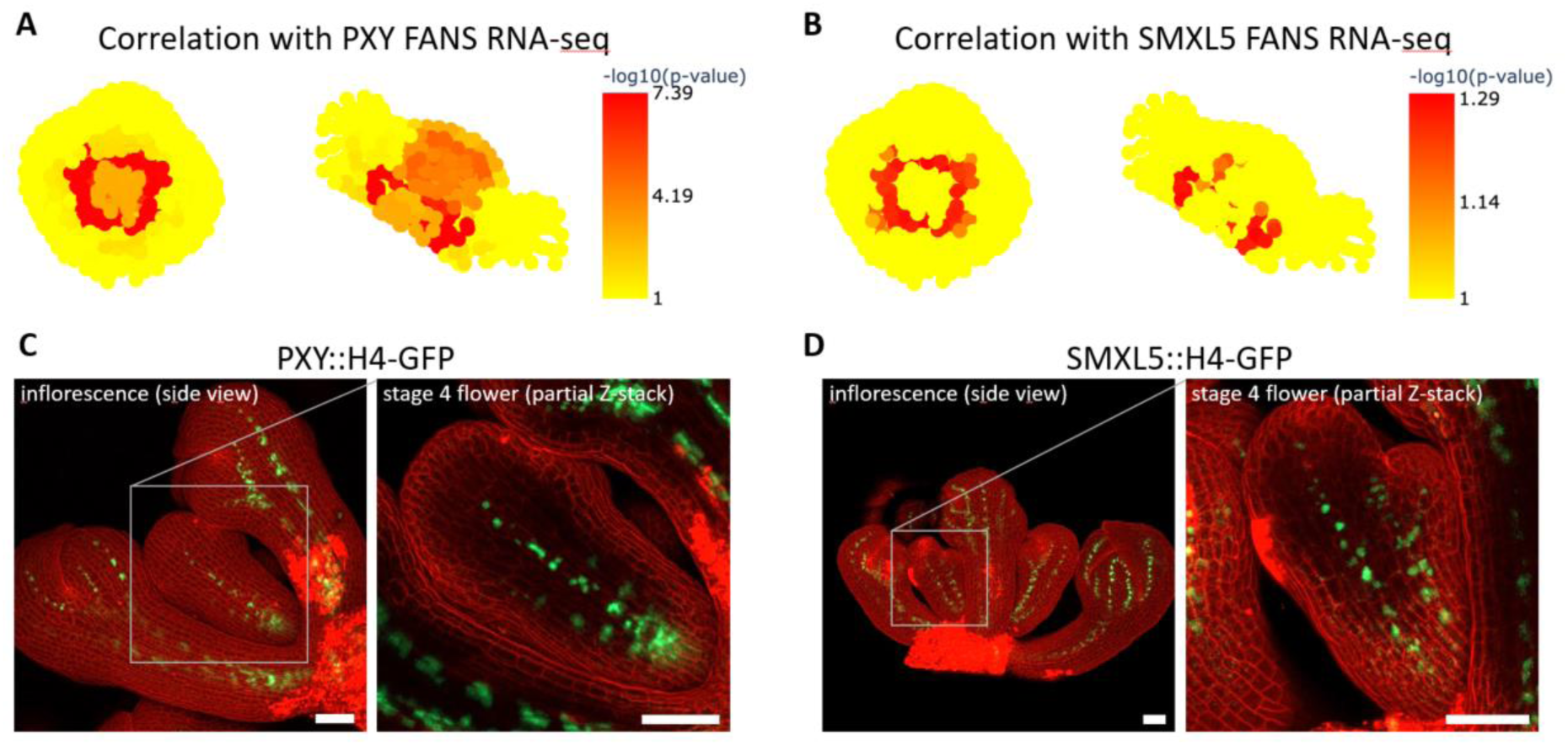
Localization of the vascular stem cells in the flower meristem. Predicted location of the vascular stem cells was calculated by the -log10 p-value of the Pearson correlation for different vascular FANS RNA-seq datasets (PXY: **A**, and SMXL5: **B**) to the reconstructed transcriptomes of each cell of the spatial map. **C** and **D** show H4-GFP expression (green) driven by the PXY and by the SMXL5 promoter, respectively. Images display side views of an inflorescence (left) and a stage 4 flower (right). For improved visualization of the GFP signal within the pedicel, the top layers of the Z-stack were removed from the orthogonal projection. Cell walls were stained using propidium iodide (red). Scale bars indicate 50 µm.

**Sup Fig 13.**
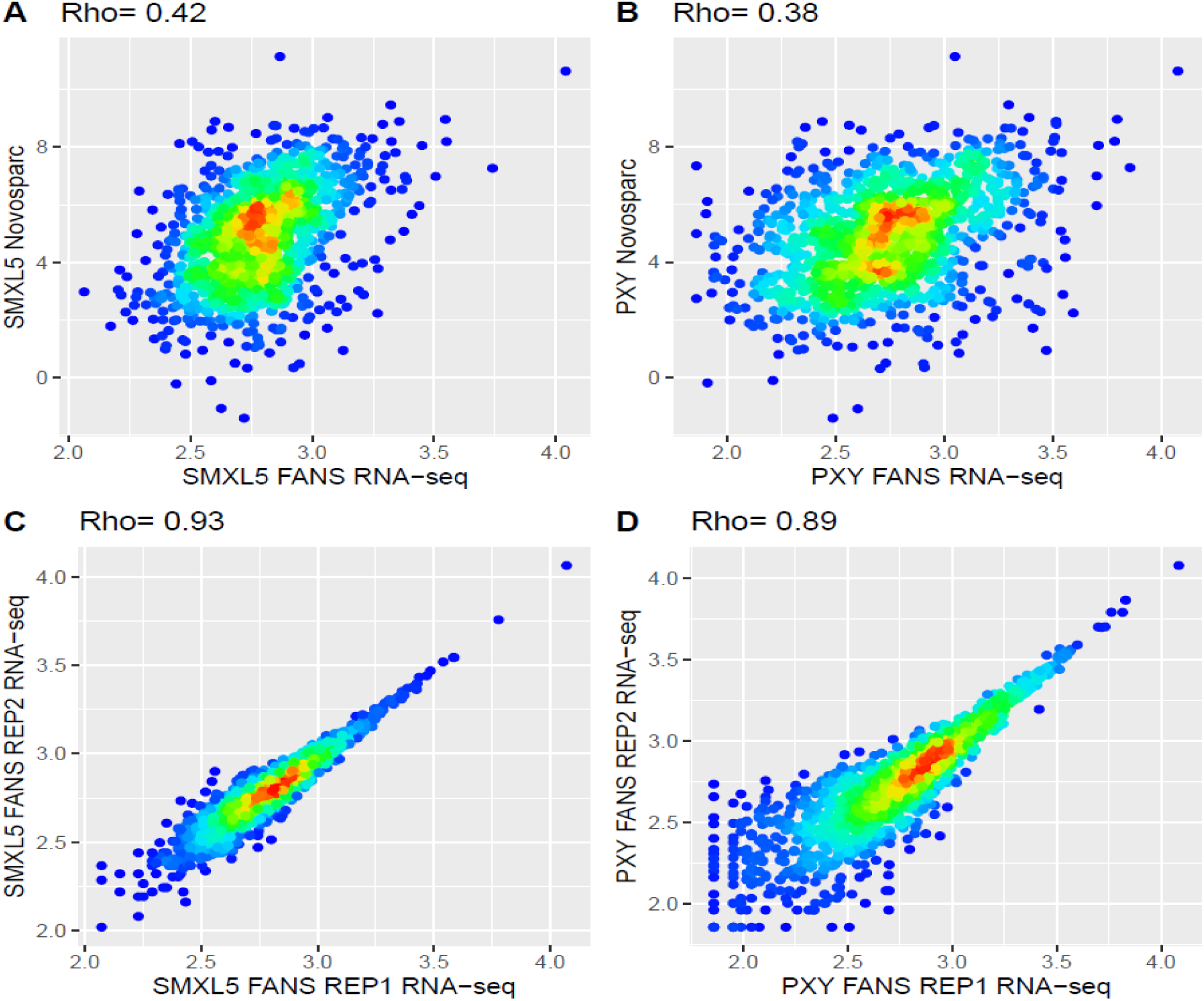
Prediction of vascular domain-specific expression. Scatterplot showing the gene expression for SMXL5 (A), and PXY (B) domain predicted by our method (y-axis) and observed by publicly available FANS bulk RNA-seq data (x-axis) when using genes with PEP value>0.13 (n=1,306). Bottom row shows the scatterplot for the gene expression of both biological FANS bulk RNA-seq replicates for SMXL5 (C), and PXY (D).

## Notes

### Competing Interest Statement

The authors have declared no competing interest.

## REFERENCES

Abdelaal, Tamim, Soufiane Mourragui, Ahmed Mahfouz, and Marcel J T Reinders. 2020. “SpaGE: Spatial Gene Enhancement Using ScRNA-Seq.” BioRxiv, January, 2020.05.08.084392. https://doi.org/10.1101/2020.05.08.084392.

Abe, Mitsutomo, Hiroshi Katsumata, Yoshibumi Komeda, and Taku Takahashi. 2003. “Regulation of Shoot Epidermal Cell Differentiation by a Pair of Homeodomain Proteins in Arabidopsis.” Development 130 (4): 635–43. https://doi.org/10.1242/dev.00292.

Bhosale, Rahul, Veronique Boudolf, Fabiola Cuevas, Ran Lu, Thomas Eekhout, Zhubing Hu, Gert Van Isterdael, et al. 2018. “A Spatiotemporal DNA Endoploidy Map of the Arabidopsis Root Reveals Roles for the Endocycle in Root Development and Stress Adaptation.” The Plant Cell 30 (10): 2330 LP – 2351. https://doi.org/10.1105/tpc.17.00983.

Bolger, Anthony M, Marc Lohse, and Bjoern Usadel. 2014. “Trimmomatic: A Flexible Trimmer for Illumina Sequence Data.” Bioinformatics 30 (15): 2114–20. https://doi.org/10.1093/bioinformatics/btu170.

Bonke, Martin, Siripong Thitamadee, Ari Pekka Mähönen, Marie-Theres Hauser, and Ykä Helariutta. 2003. “APL Regulates Vascular Tissue Identity in Arabidopsis.” Nature 426 (6963): 181–86. https://doi.org/10.1038/nature02100.

Bravo González-Blas, Carmen, Xiao-Jiang Quan, Ramon Duran-Romaña, Ibrahim Ihsan Taskiran, Duygu Koldere, Kristofer Davie, Valerie Christiaens, et al. 2020. “Identification of Genomic Enhancers through Spatial Integration of Single-cell Transcriptomics and Epigenomics.” Molecular Systems Biology 16 (5). https://doi.org/10.15252/msb.20209438.

Brewer, Philip B, Paul A Howles, Kristen Dorian, Megan E Griffith, Tetsuya Ishida, Ruth N Kaplan-Levy, Aydin Kilinc, and David R Smyth. 2004. “PETAL LOSS, a Trihelix Transcription Factor Gene, Regulates Perianth Architecture in the Arabidopsis Flower.” Development 131 (16): 4035–45. https://doi.org/10.1242/dev.01279.

Cheng, Chia-Yi, Vivek Krishnakumar, Agnes P Chan, Françoise Thibaud-Nissen, Seth Schobel, and Christopher D Town. 2017. “Araport11: A Complete Reannotation of the Arabidopsis Thaliana Reference Genome.” The Plant Journal 89 (4): 789–804. https://doi.org/https://doi.org/10.1111/tpj.13415.

Chuang, Chiou Fen, Mark P. Running, Robert W. Williams, and Elliot M. Meyerowitz. 1999. “The PERIANTHIA Gene Encodes a BZIP Protein Involved in the Determination of Floral Organ Number in Arabidopsis Thaliana.” Genes and Development 13 (3): 334–44. https://doi.org/10.1101/gad.13.3.334.

Deal, Roger B, and Steven Henikoff. 2011. “The INTACT Method for Cell Type–Specific Gene Expression and Chromatin Profiling in Arabidopsis Thaliana.” Nature Protocols 6 (1): 56–68. https://doi.org/10.1038/nprot.2010.175.

Denyer, Tom, Xiaoli Ma, Simon Klesen, Emanuele Scacchi, Kay Nieselt, and Marja C.P. Timmermans. 2019. “Spatiotemporal Developmental Trajectories in the Arabidopsis Root Revealed Using High-Throughput Single-Cell RNA Sequencing.” Developmental Cell 48 (6): 840–852.e5. https://doi.org/10.1016/j.devcel.2019.02.022.

Dobin, Alexander, Carrie A Davis, Felix Schlesinger, Jorg Drenkow, Chris Zaleski, Sonali Jha, Philippe Batut, Mark Chaisson, and Thomas R Gingeras. 2013. “STAR: Ultrafast Universal RNA-Seq Aligner.” Bioinformatics 29 (1): 15–21. https://doi.org/10.1093/bioinformatics/bts635.

Duncan, Susan, Tjelvar S G Olsson, Matthew Hartley, Caroline Dean, and Stefanie Rosa. 2016. “A Method for Detecting Single MRNA Molecules in Arabidopsis Thaliana.” Plant Methods 12 (1): 13. https://doi.org/10.1186/s13007-016-0114-x.

Giacomello, Stefania, Fredrik Salmén, Barbara K. Terebieniec, Sanja Vickovic, José Fernandez Navarro, Andrey Alexeyenko, Johan Reimegård, et al. 2017. “Spatially Resolved Transcriptome Profiling in Model Plant Species.” Nature Plants 3 (May). https://doi.org/10.1038/nplants.2017.61.

Hassan, Hala, Ben Scheres, and Ikram Blilou. 2010. “JACKDAW Controls Epidermal Patterning in the Arabidopsis Root Meristem through a Non-Cell-Autonomous Mechanism.” Development 137 (9): 1523–29. https://doi.org/10.1242/dev.048777.

Hernandez-Lagana, Elvira, Gabriella Mosca, Ethel Mendocilla-Sato, Nuno Pires, Anja Frey, Alejandro Giraldo-Fonseca, Caroline Michaud, et al. 2021. “Organ Geometry Channels Reproductive Cell Fate in the Arabidopsis Ovule Primordium.” Edited by Sheila McCormick. ELife 10: e66031. https://doi.org/10.7554/eLife.66031.

Jack, Thomas, Laura L Brockman, and Elliot M Meyerowitz. 1992. “The Homeotic Gene APETALA3 of Arabidopsis Thaliana Encodes a MADS Box and Is Expressed in Petals and Stamens.” Cell 68 (4): 683–97. https://doi.org/https://doi.org/10.1016/0092-8674(92)90144-2.

Jensen, Anders B, Dora Raventos, and John Mundy. 2002. “Fusion Genetic Analysis of Jasmonate-Signalling Mutants in Arabidopsis.” The Plant Journal 29 (5): 595–606. https://doi.org/https://doi.org/10.1046/j.0960-7412.2001.01241.x.

Karaiskos, Nikos, Philipp Wahle, Jonathan Alles, Anastasiya Boltengagen, Salah Ayoub, Claudia Kipar, Christine Kocks, Nikolaus Rajewsky, and Robert P Zinzen. 2017. “The *Drosophila* Embryo at Single-Cell Transcriptome Resolution.” Science 358 (6360): 194 LP – 199. https://doi.org/10.1126/science.aan3235.

Kaufmann, Kerstin, Frank Wellmer, Jose M. Muiñ, Thilia Ferner, Samuel E. Wuest, Vijaya Kumar, Antonio Serrano-Mislata, et al. 2010. “Orchestration of Floral Initiation by APETALA1.” Science 328 (5974): 85–89. https://doi.org/10.1126/science.1185244.

Klepikova, Anna V, Artem S Kasianov, Evgeny S Gerasimov, Maria D Logacheva, and Aleksey A Penin. 2016. “A High Resolution Map of the Arabidopsis Thaliana Developmental Transcriptome Based on RNA-Seq Profiling.” The Plant Journal 88 (6): 1058–70. https://doi.org/https://doi.org/10.1111/tpj.13312.

Korsunsky, Ilya, Nghia Millard, Jean Fan, Kamil Slowikowski, Fan Zhang, Kevin Wei, Yuriy Baglaenko, Michael Brenner, Po-ru Loh, and Soumya Raychaudhuri. 2019. “Fast, Sensitive and Accurate Integration of Single-Cell Data with Harmony.” Nature Methods 16 (12): 1289–96. https://doi.org/10.1038/s41592-019-0619-0.

Kunst, L, J E Klenz, J Martinez-Zapater, and G W Haughn. 1989. “AP2 Gene Determines the Identity of Perianth Organs in Flowers of Arabidopsis Thaliana.” The Plant Cell 1 (12): 1195–1208. https://doi.org/10.1105/tpc.1.12.1195.

Larkin, Robert M, Giovanni Stefano, Michael E Ruckle, Andrea K Stavoe, Christopher A Sinkler, Federica Brandizzi, Carolyn M Malmstrom, and Katherine W Osteryoung. 2016. “*REDUCED CHLOROPLAST COVERAGE* Genes from *Arabidopsis Thaliana* Help to Establish the Size of the Chloroplast Compartment.” Proceedings of the National Academy of Sciences 113 (8): E1116 LP-E1125. https://doi.org/10.1073/pnas.1515741113.

Liao, Yang, Gordon K Smyth, and Wei Shi. 2014. “FeatureCounts: An Efficient General Purpose Program for Assigning Sequence Reads to Genomic Features.” Bioinformatics 30 (7): 923–30. https://doi.org/10.1093/bioinformatics/btt656.

Lopez, Romain, Jeffrey Regier, Michael B Cole, Michael I Jordan, and Nir Yosef. 2018. “Deep Generative Modeling for Single-Cell Transcriptomics.” Nature Methods 15 (12): 1053–58. https://doi.org/10.1038/s41592-018-0229-2.

Love, Michael I, Wolfgang Huber, and Simon Anders. 2014. “Moderated Estimation of Fold Change and Dispersion for RNA-Seq Data with DESeq2.” Genome Biology 15 (12): 550. https://doi.org/10.1186/s13059-014-0550-8.

Mantegazza, Otho, Veronica Gregis, Marta Adelina Mendes, Piero Morandini, Márcio Alves-Ferreira, Camila M Patreze, Sarah M Nardeli, Martin M Kater, and Lucia Colombo. 2014. “Analysis of the Arabidopsis REM Gene Family Predicts Functions during Flower Development.” Annals of Botany 114 (7): 1507–15. https://doi.org/10.1093/aob/mcu124.

Marx, Vivien. 2021. “Method of the Year: Spatially Resolved Transcriptomics.” Nature Methods 18 (1): 9–14. https://doi.org/10.1038/s41592-020-01033-y.

Mizukami, Yukiko, and Robert L Fischer. 2000. “Plant Organ Size Control: *AINTEGUMENTA* Regulates Growth and Cell Numbers during Organogenesis.” Proceedings of the National Academy of Sciences 97 (2): 942 LP – 947. https://doi.org/10.1073/pnas.97.2.942.

Nitzan, Mor, Nikos Karaiskos, Nir Friedman, and Nikolaus Rajewsky. 2019. “Gene Expression Cartography.” Nature 576 (7785): 132–37. https://doi.org/10.1038/s41586-019-1773-3.

Ó’Maoiléidigh, Diarmuid S, Samuel E Wuest, Liina Rae, Andrea Raganelli, Patrick T Ryan, Kamila Kwaśniewska, Pradeep Das, et al. 2013. “Control of Reproductive Floral Organ Identity Specification in Arabidopsis by the C Function Regulator AGAMOUS .” The Plant Cell 25 (7): 2482–2503. https://doi.org/10.1105/tpc.113.113209.

Pajoro, A., P. Madrigal, J.M. Muiño, J.T. Matus, J. Jin, M.A. Mecchia, J.M. Debernardi, et al. 2014. “Dynamics of Chromatin Accessibility and Gene Regulation by MADS-Domain Transcription Factors in Flower Development.” Genome Biology 15 (3). https://doi.org/10.1186/gb-2014-15-3-r41.

Pesch, Martina, and Martin Hülskamp. 2011. “Role of TRIPTYCHON in Trichome Patterning in Arabidopsis.” BMC Plant Biology 11 (1): 130. https://doi.org/10.1186/1471-2229-11-130.

Refahi, Yassin, Argyris Zardilis, Gaël Michelin, Raymond Wightman, Bruno Leggio, Jonathan Legrand, Emmanuel Faure, et al. 2021. “A Multiscale Analysis of Early Flower Development in Arabidopsis Provides an Integrated View of Molecular Regulation and Growth Control.” Developmental Cell 56 (4): 540–556.e8. https://doi.org/10.1016/j.devcel.2021.01.019.

Samach, Alon, Jennifer E Klenz, Susanne E Kohalmi, Eddy Risseeuw, George W Haughn, and William L Crosby. 1999. “The UNUSUAL FLORAL ORGANS Gene of Arabidopsis Thaliana Is an F-Box Protein Required for Normal Patterning and Growth in the Floral Meristem.” The Plant Journal 20 (4): 433–45. https://doi.org/https://doi.org/10.1046/j.1365-313x.1999.00617.x.

Sanchez, Pablo, Lilian Nehlin, and Thomas Greb. 2012. “From Thin to Thick: Major Transitions during Stem Development.” Trends in Plant Science 17 (2): 113–21. https://doi.org/10.1016/j.tplants.2011.11.004.

Satija, Rahul, Jeffrey A Farrell, David Gennert, Alexander F Schier, and Aviv Regev. 2015. “Spatial Reconstruction of Single-Cell Gene Expression Data.” Nature Biotechnology 33 (5): 495– 502. https://doi.org/10.1038/nbt.3192.

Sessions, Allen, Detlef Weigel, and Martin F Yanofsky. 1999. “The Arabidopsis Thaliana MERISTEM LAYER 1 Promoter Specifies Epidermal Expression in Meristems and Young Primordia.” The Plant Journal 20 (2): 259–63. https://doi.org/https://doi.org/10.1046/j.1365-313x.1999.00594.x.

Shi, Dongbo, Virginie Jouannet, Javier Agustí, Verena Kaul, Victor Levitsky, Pablo Sanchez, Victoria V Mironova, and Thomas Greb. 2021. “Tissue-Specific Transcriptome Profiling of the Arabidopsis Inflorescence Stem Reveals Local Cellular Signatures.” The Plant Cell 33 (2): 200–223. https://doi.org/10.1093/plcell/koaa019.

Smyth, David R., John L. Bowman, and Elliot M. Meyerowitz. 1990. “Early Flower Development in Arabidopsis.” Plant Cell 2 (8): 755–67. https://doi.org/10.1105/tpc.2.8.755.

Solanki, Shyam, Gazala Ameen, Jin Zhao, Jordan Flaten, Pawel Borowicz, and Robert S Brueggeman. 2020. “Visualization of Spatial Gene Expression in Plants by Modified RNAscope Fluorescent in Situ Hybridization.” Plant Methods 16 (1): 71. https://doi.org/10.1186/s13007-020-00614-4.

Stuart, Tim, Andrew Butler, Paul Hoffman, Christoph Hafemeister, Efthymia Papalexi, William M Mauck III, Yuhan Hao, Marlon Stoeckius, Peter Smibert, and Rahul Satija. 2019. “Comprehensive Integration of Single-Cell Data.” Cell 177 (7): 1888–1902.e21. https://doi.org/10.1016/j.cell.2019.05.031.

Sunaga-Franze, Daniele Y, Jose M Muino, Caroline Braeuning, Xiaocai Xu, Minglei Zong, Cezary Smaczniak, Wenhao Yan, et al. 2021. “Single-Nuclei RNA-Sequencing of Plant Tissues.” BioRxiv, January, 2020.11.14.382812. https://doi.org/10.1101/2020.11.14.382812.

Tian, Caihuan, Ying Wang, Haopeng Yu, Jun He, Jin Wang, Bihai Shi, Qingwei Du, Nicholas J Provart, Elliot M Meyerowitz, and Yuling Jiao. 2019. “A Gene Expression Map of Shoot Domains Reveals Regulatory Mechanisms.” Nature Communications 10 (1): 141. https://doi.org/10.1038/s41467-018-08083-z.

Uchida, Naoyuki, Jin Suk Lee, Robin J Horst, Hung-Hsueh Lai, Ryoko Kajita, Tatsuo Kakimoto, Masao Tasaka, and Keiko U Torii. 2012. “Regulation of Inflorescence Architecture by Intertissue Layer Ligand–Receptor Communication between Endodermis and Phloem.” Proceedings of the National Academy of Sciences 109 (16): 6337 LP – 6342. https://doi.org/10.1073/pnas.1117537109.

Valuchova, Sona, Pavlina Mikulkova, Jana Pecinkova, Jana Klimova, Michal Krumnikl, Petr Bainar, Stefan Heckmann, Pavel Tomancak, and Karel Riha. 2020. “Imaging Plant Germline Differentiation within Arabidopsis Flowers by Light Sheet Microscopy.” ELife 9: e52546. https://doi.org/10.7554/eLife.52546.

Vijayan, Athul, Rachele Tofanelli, Sören Strauss, Lorenzo Cerrone, Adrian Wolny, Joanna Strohmeier, Anna Kreshuk, Fred A Hamprecht, Richard S Smith, and Kay Schneitz. 2021. “A Digital 3D Reference Atlas Reveals Cellular Growth Patterns Shaping the Arabidopsis Ovule.” Edited by Sheila McCormick, Christian S Hardtke, Sheila McCormick, and Dolf Weijers. ELife 10: e63262. https://doi.org/10.7554/eLife.63262.

Wang, Shucai, Su-Hwan Kwak, Qingning Zeng, Brian E Ellis, Xiao-Ya Chen, John Schiefelbein, and Jin-Gui Chen. 2007. “TRICHOMELESS1 Regulates Trichome Patterning by Suppressing GLABRA1 in Arabidopsis.” Development 134 (21): 3873–82. https://doi.org/10.1242/dev.009597.

Waylen, Lisa N, Hieu T Nim, Luciano G Martelotto, and Mirana Ramialison. 2020. “From Whole-Mount to Single-Cell Spatial Assessment of Gene Expression in 3D.” Communications Biology 3 (1): 602. https://doi.org/10.1038/s42003-020-01341-1.

Welch, Joshua D, Velina Kozareva, Ashley Ferreira, Charles Vanderburg, Carly Martin, and Evan Z Macosko. 2019. “Single-Cell Multi-Omic Integration Compares and Contrasts Features of Brain Cell Identity.” Cell 177 (7): 1873–1887.e17. https://doi.org/https://doi.org/10.1016/j.cell.2019.05.006.

Wolny, Adrian, Lorenzo Cerrone, Athul Vijayan, Rachele Tofanelli, Amaya Vilches Barro, Marion Louveaux, Christian Wenzl, et al. 2020. “Accurate and Versatile 3D Segmentation of Plant Tissues at Cellular Resolution.” ELife 9 (July). https://doi.org/10.7554/elife.57613.

Wuest, Samuel E., Diarmuid S. O’Maoileidigh, Liina Rae, Kamila Kwasniewska, Andrea Raganelli, Katarzyna Hanczaryk, Amanda J. Lohan, Brendan Loftus, Emmanuelle Graciet, and Frank Wellmer. 2012. “Molecular Basis for the Specification of Floral Organs by APETALA3 and PISTILLATA.” Proceedings of the National Academy of Sciences of the United States of America 109 (33): 13452–57. https://doi.org/10.1073/pnas.1207075109.

Xu, Xiaocai, Cezary Smaczniak, Jose M Muino, and Kerstin Kaufmann. 2021. “Cell Identity Specification in Plants: Lessons from Flower Development.” *Journal of Experimental Botany*, April. https://doi.org/10.1093/jxb/erab110.

Yadav, Ram Kishor, Montreh Tavakkoli, Mingtang Xie, Thomas Girke, and G Venugopala Reddy. 2014. “A High-Resolution Gene Expression Map of the Arabidopsis Shoot Meristem Stem Cell Niche.” Development 141 (13): 2735–44. https://doi.org/10.1242/dev.106104.

Yang, Weibing, Raymond Wightman, and Elliot M Meyerowitz. 2017. “Cell Cycle Control by Nuclear Sequestration of CDC20 and CDH1 MRNA in Plant Stem Cells.” Molecular Cell 68 (6): 1108–1119.e3. https://doi.org/10.1016/j.molcel.2017.11.008.

